# GTF2I dosage regulates neuronal differentiation and social behavior in 7q11.23 neurodevelopmental disorders

**DOI:** 10.1101/2022.10.10.511434

**Authors:** Alejandro Lopez-Tobon, Reinald Shyti, Carlo Emanuele Villa, Cristina Cheroni, Patricio Fuentes-Bravo, Sebastiano Trattaro, Nicolò Caporale, Flavia Troglio, Erika Tenderini, Marija Mihailovich, Adrianos Skaros, William T. Gibson, Alessandro Cuomo, Tiziana Bonaldi, Ciro Mercurio, Mario Varasi, Lucy Osborne, Giuseppe Testa

## Abstract

Copy number variations at 7q11.23 cause neurodevelopmental disorders with shared and opposite manifestations. Deletion causes Williams-Beuren syndrome (WBS), while duplication causes 7q11.23 microduplication syndrome (7Dup). Converging evidence indicates *GTF2I*, from the 7q11.23 locus, is a key mediator of the cognitive-behavioral phenotypes associated with WBS and 7Dup. Here we integrate molecular profiling of patient-derived cortical organoids (COs) and transgenic mouse models to dissect 7q11.23 disease mechanisms. Proteomic and transcriptomic profiling of COs revealed opposite dynamics of neural progenitor proliferation and transcriptional imbalances, leading to precocious excitatory neuron production in 7Dup. The accelerated excitatory neuron production in 7Dup COs could be rescued by *GTF2I* knockdown. Transgenic mice with *Gtf2i* duplication recapitulated early neuronal differentiation defects and ASD-like behaviors. Remarkably, inhibition of LSD1, a downstream effector of *GTF2I*, was sufficient to rescue ASD-like phenotypes. We propose that the GTF2I-LSD1 axis constitutes a molecular pathway amenable to therapeutic intervention.

## Introduction

Copy number variations (CNVs) at the 7q11.23 locus spanning 26-28 genes cause two rare neurodevelopmental disorders featuring shared and opposite multisystemic phenotypes. Hemi deletion causes Williams-Beuren Syndrome (WBS; OMIM (Online Mendelian Inheritance in Man) 194050), featuring intellectual disability coupled with hypersociability, anxiety and comparatively well-preserved language abilities (Pober, 2010). Meanwhile, 7q11.23 hemi duplication (7Dup; OMIM 609757) shares intellectual disability and anxiety, with a contrasting impairment in expressive language and a high incidence of ASD (Sanders et al., 2011; Van der Aa et al., 2009). Due to their unique combination of symmetrically opposite neuropsychiatric manifestations and genetic lesions, 7q11.23 CNV syndromes offer unprecedented opportunities to dissect genes and their dosage-vulnerable circuits involved in social behavior and language competence (López-Tobón et al., 2020). Likewise, recent initiatives in large human samples have linked genome dosage sensitive regions with specific disorders, revealing CNVs properties and identifying thousands of highly dosage sensitive genes (Collins et al., 2022)

Convergent evidence from mouse and human studies, including WBS individuals carrying atypical deletions, have pointed to General Transcription factor I (*GTF2I*) as a key driver of the 7q11.23 CNV cognitive-behavioral manifestations (Antonell et al., 2010; Crespi and Hurd, 2014), however its mechanistic underpinnings and specific roles during cortical development and in behavioral alterations remains elusive. *GTF2I* is a signal-induced transcription factor and its target genes are involved in a variety of neuronal functions including axon guidance, calcium signaling, cell cycle and maturation of GABA-ergic interneurons (Chimge et al., 2008; Poitras et al., 2010; Roy, 2017). Previously, we defined the *GTF2I* dose-dependent transcriptional dysregulation at the pluripotent state, and traced its amplification upon differentiation into neural progenitors. Critically, we discovered that *GTF2I* represses transcription by binding the histone lysine demethylase 1 (LSD1 or KDM1A), whereas chemical and genetic LSD1 inhibition rescued the aberrant transcriptional repression brought about by increased GTF2I dosage in 7Dup lineages, affecting expression of key genes involved in intellectual disability and autism and underlining the potential of LSD1 inhibition for therapeutic intervention (Adamo et al., 2015).

The use of patient-derived pluripotent stem cell models, in particular brain organoids in combination with single cell transcriptomics, is a powerful platform to model neuropsychiatric disorders with unprecedented accuracy (Mariani et al., 2015; Paulsen et al., 2022; Velmeshev et al., 2019). Its application to study processes that reach unparalleled complexity in humans, such as the genetic circuits regulating cortical development and behavior, holds enormous potential to uncover hubs of genetic regulation susceptible to impairment during development. As a testament to the potential of these approaches, we and others recently showed that autism spectrum disorder (ASD)-risk genes’ pathogenic impact converge on alterations of neuronal differentiation dynamics and neurodevelopmental trajectories (Paulsen et al., 2022; Villa et al., 2022).

Here we undertook a systematic dissection of the impact of 7q11.23 CNVs in cortical development, juxtaposing WBS and 7Dup transcriptional and phenotypic landscapes across multiple individuals, key developmental time windows and experimental models. Our results, from integrating cortical organoids (CO) with single cell transcriptomics, revealed symmetrically opposite defects in progenitor proliferation and neuronal maturation dynamics as a result of 7q11.23 gene dosage imbalances. Remarkably, changes in *GTF2I* dosage alone, either in COs or transgenic mice, were sufficient to phenocopy key neuronal differentiation alterations observed in COs and caused ASD-like behaviors that could be rescued by inhibition of the downstream *GTF2I* effector, LSD1.

## Results

### 7q11.23 reciprocal CNVs causes progenitor proliferation and neuronal differentiation imbalances during corticogenesis in cortical organoids

We generated patient-derived COs that recapitulate early to mid-fetal corticogenesis (Cheroni et al., 2022; Paşca et al., 2015; Yoon et al., 2019) from 13 iPSC lines reprogrammed from 8 affected individuals and 5 healthy controls (4-WBS, 4-7Dup and 5 CTL, **Figure 1A**), including, respectively, 1 clone per individual following the design that we previously benchmarked (Germain and Testa, 2017), in order to avoid artificial inflation of spurious differential expression due to donors‘ genetic backgrounds. Array comparative genomic hybridization (aCGH) confirmed that the 7q11.23 CNV is preserved upon reprogramming (**Figure S1A**). Likewise, the transcriptional readout of the locus dosage imbalance was confirmed by unattended clustering of the expression levels of the 7q11.23 genes in 7Dup, WBS and CTL COs at day 18, 50 and 100 of differentiation (**Figure S1B**). Interestingly, the magnitude of the average locus fold-change is not constant, but peaks at day 18 of differentiation (**Figure S1C**), pointing to developmental windows of differential dosage sensitivity in terms of transcriptional readout.

**Figure 1.**
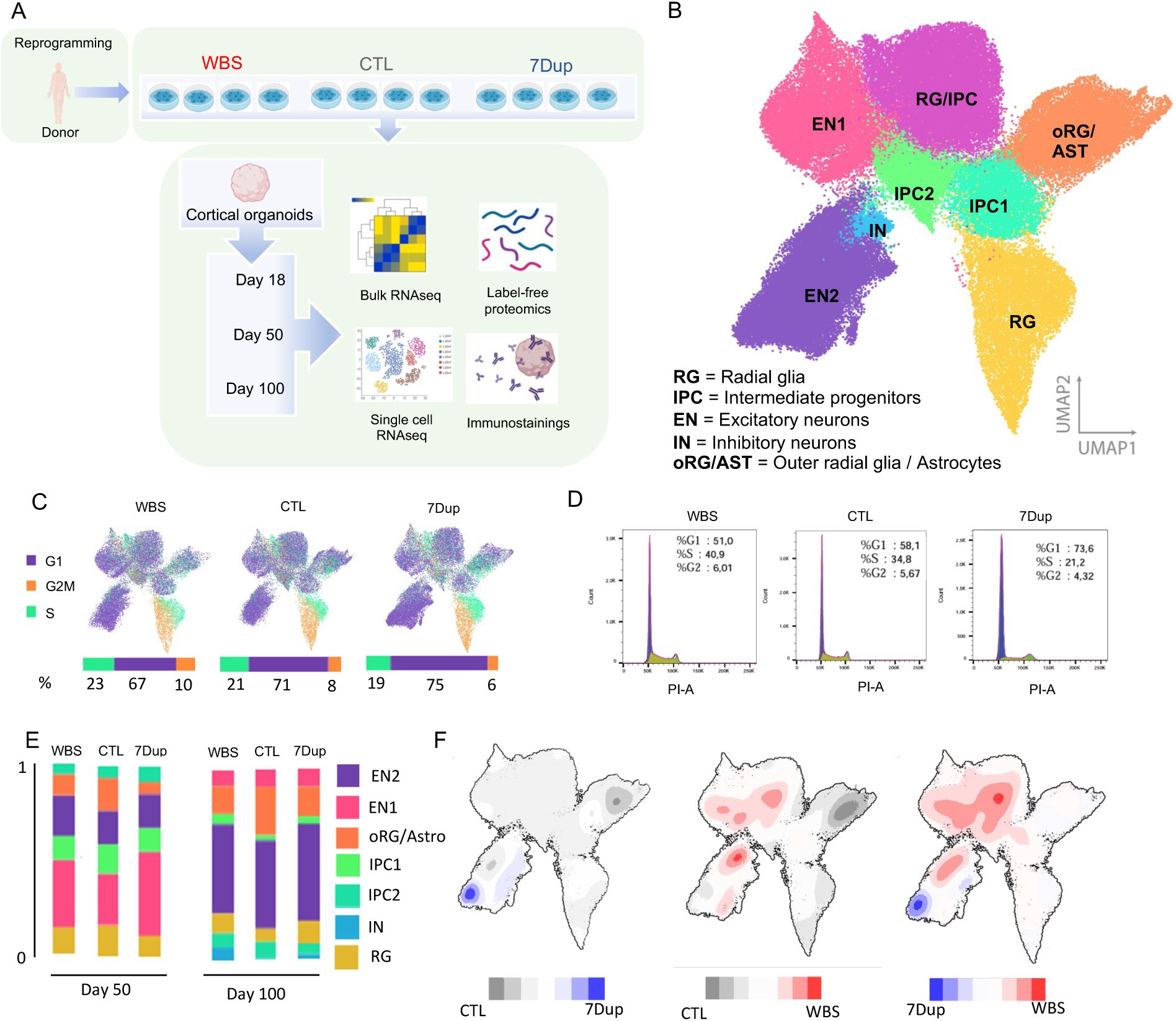
7q11.23 CNVs cause aberrations in cell cycle and neuronal population composition in COs. **(A)** Experimental design: 13 reprogrammed iPSC lines from 13 different donors (4 - WBS, 5- CTL and 4- 7Dup) were differentiated into COs and profiled for bulk transcriptomics, scRNAseq and whole proteome at the specified timepoints. **(B)** Uniform manifold approximation and projection (UMAP) from (n = 97108) single cell transcriptomes. Leiden clustering algorithm identified 8 distinct cell cluster (RG, RG/IPC, IPC1-2, EN1-2, IN, oRG/AST), composed by the following populations: Radial Glia (RG), Intermediate Progenitors (IPC), Excitatory Neurons (EN), Inhibitory Neurons (IN), Outer radial glia / Astrocytes (oRG/AST). **(C)** Percentage of total amount of cells in each cell cycle phase and distribution in UMAP stratified by condition at day 50. **(D)** Propidium iodine flow cytometry quantification from day 18 organoids (N = 4 - WBS, 4- CTL and 4- 7Dup), pool of 3 organoids per genotype. **(E)** Proportion of cells from each genotype in each cell cluster in COs at 50 days and 100 days (3 pooled individual organoids per genotype) **(F)** Density plots illustrating differences in cell abundance in UMAP discriminated by condition; Red = high abundance in WBS, Blue = high abundance in 7Dup, Grey = high abundance in control.

To investigate the impact of 7q11.23 CNVs on corticogenesis, we profiled COs at 3 time-points (**Figure 1A**) that we previously showed have a selective enrichment in ventricular radial glia (vRG; day 18), intermediate progenitors cells (IPC; day 50), outer radial glia and excitatory neurons (oRG; day 100), a progenitor population essential for cortical expansion in humans (López-Tobón et al., 2019; Pollen et al., 2015). Droplet-based transcriptional profiling of ∼100.000 single cells from patient-derived COs (day 50 and 100) identified 8 cell population clusters encompassing radial glial cells (RG), IPC, astrocytes (AST) and excitatory neurons (EN) (**Figure 1B**). Population annotation was achieved by tracing the expression pattern of known cortical population markers (**Figure S2A**), unattended determination of cluster-specific highly expressed genes (**Figure S2B**) and overlap with public fetal cortical single cell datasets (**Figure S2C,D**). We confirmed that control lines showed reproducible proportions of all identified populations (**Figure S2E**), as previously shown for this CO protocol (Gordon et al., 2021; Yoon et al., 2019), and proceeded to ascertain that each line independently reproduced stereotypical developmental trajectories from radial glia progenitors to mature excitatory neurons and astrocytes (**Figure S2F**).

Quantification of the cell cycle phase distribution between WBS and 7Dup COs at day 50 revealed symmetrically opposite proportions of cells in S – G2M phases (WBS % = 33, CTL % = 29 and 7Dup % = 25) (**Figure 1C**). This cycling asymmetry was already present at day 18 of differentiation, as indicated by the observed shift in cell cycle proportions (**Figure 1D**). Consistently, comparison of KI67^+^ cells at day 50 confirmed an increase of proliferating cells in WBS and a decrease in 7Dup (**Figure S3A**), defining a symmetrically opposite imbalance in cell proliferation across 7q11.23 genotypes during early corticogenesis.

To probe how these divergent proliferative potential impacts different cortical neuronal populations, we immunostained and quantified population markers of early to mid-corticogenesis. We found a consistent increase in the proportion of PAX6^+^ progenitors and TBR2^+^ intermediate progenitors in WBS compared with 7Dup COs (**Figure S3D**) while, conversely, postmitotic neurons positive for Layer V-VI markers (TBR1^+^ BCL11B^+^) were increased in 7Dup COs compared with WBS (**Figure S3E-G**). Furthermore, we examined the proportion of cell populations across conditions through days 50-100, finding both common and symmetrically opposite changes between the two conditions. Specifically, the changes in cell population frequencies common to both WBS and 7Dup COs included a 2-fold reduction in the number of oRG/Astro cells, a 1.5-fold increase in the total number of excitatory neurons and the appearance of a population rich in ventral telencephalon and interneuron (IN) markers (DLX1/2) (**Figure 1E**). Symmetrically opposite alterations were instead apparent within cell clusters upon detailed mapping by density plot comparing either condition to the control and to each other (**Figure 1F**), revealing mirroring frequencies of excitatory neurons (increased in 7Dup and decreased in WBS) alongside a WBS-specific increase in RG/IPCs. These results establish that gene dosage imbalances within the 7q11.23 locus impact corticogenesis by altering the dynamics of progenitor proliferation and the subsequent output of neuronal populations.

### Transcriptomic and proteomic profiling of 7q11.23 CNV COs confirm symmetrically opposite alterations in proliferative and neurogenic programs in WBS and 7Dup

In order to identify the gene circuitries involved in the population imbalances caused by 7q11.23 CNVs, we performed bulk transcriptomic profiling of COs at day 18, 50 and 100, interrogating the effect of the opposite gene dosages in developmental gene expression networks through a differential expression analysis (DEA) comparing 7Dup and WBS COs.

First, we focused on the transcriptional changes that persisted throughout differentiation by differential analysis of the complete CO cohort, specifying the differentiation stage as co-variable in the model. This analysis revealed 164 differentially expressed genes (DEGs) (FDR < 0.1, LogFC > 1 as absolute value) (**Figure 2A**). 7Dup COs showed an upregulation of the forebrain development regulator *(FOXG1)*, as well as glutamatergic lineage drivers, including (*NEUROD6*) and (*TBR1*) (**Figure 2B**), which regulate neuronal fate specification and differentiation during cortical development and are recurrently disrupted in ASD (Mariani et al., 2015; O’Roak et al., 2012; Tutukova et al., 2021). In addition, 7Dup COs showed an upregulation of members of the protocadherin family (i.e., *PCDHGA3, PCDHGA4, PCDHB5, PCDHB15, PCDHGB3*) (**Figure 2B**) known to regulate neuronal connectivity, extracellular matrix (ECM) remodeling and synapse organization (Takeichi, 2007). Conversely, WBS organoids showed an upregulation of TFs involved in anterior-lateral brain patterning (*PHOX2A, PHOX2B*) (Roome et al., 2020) as well as the G-coupled protein receptor (*GPR50*), known for its association with bipolar affective disorder, major depressive disorder and schizophrenia (Thomson et al., 2005).

**Figure 2.**
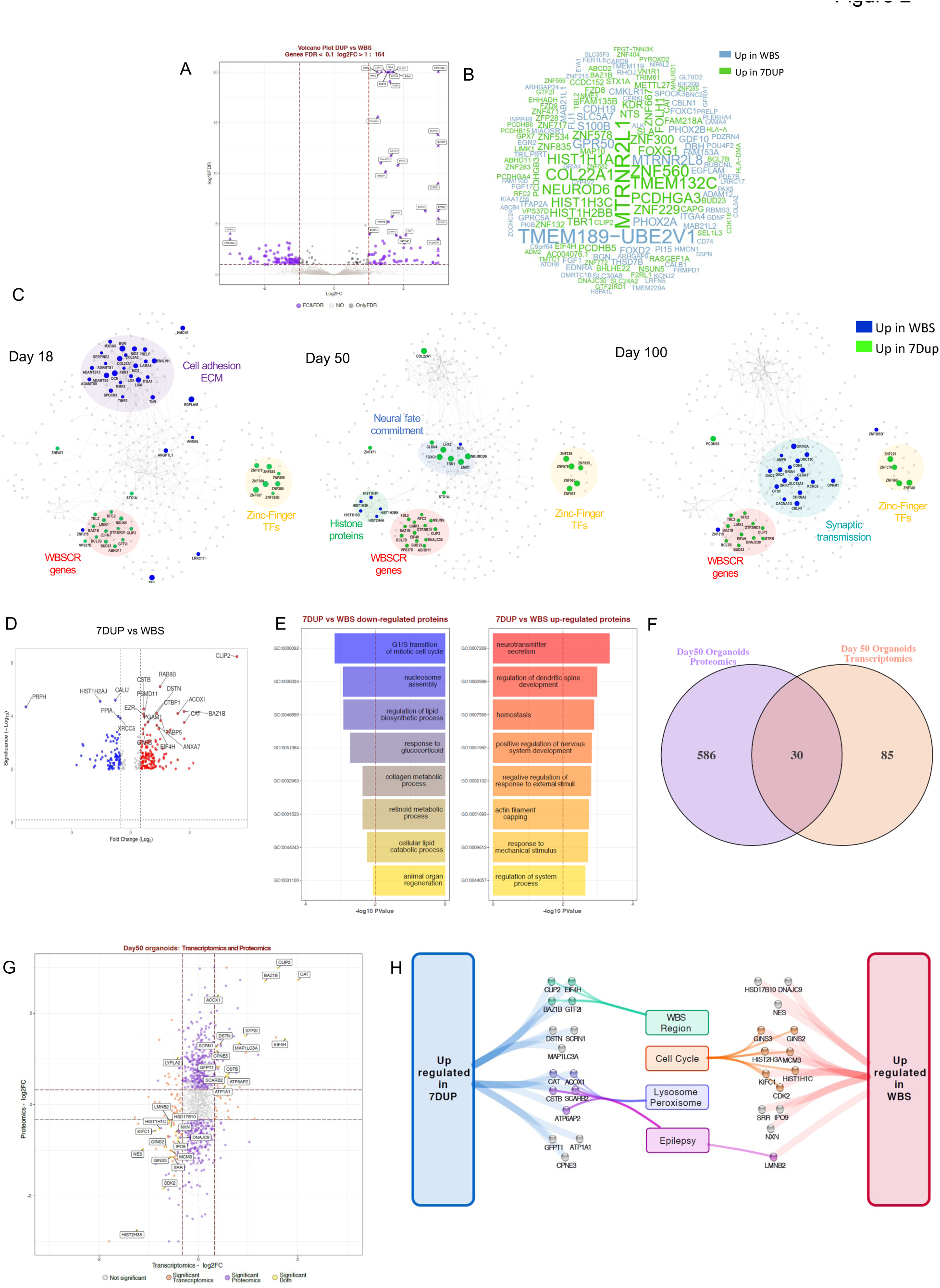
Transcriptomic and proteomic profiling of 7q11.23-CNVs COs reveal imbalances in proliferative *vis-a-vis* neurogenic programs. **(A)** Volcano plots illustrating the differential expression analysis for 7Dup vs WBS on the complete organoid cohort profiled by bulkRNASeq (33 samples: 3 WBS, 4 CTL and 3 7Dup for Day18 organoids; 4 WBS, 4 CTL and 4 7Dup for Day50 organoids; 3 WBS, 5 CTL and 3 7Dup for Day100 organoids). Results are reported for the pool of tested genes as –log10 false discovery rate (FDR) and log2 fold change (FC). Genes identified as significantly modulated (FDR < 0.1 and absolute log2FC > 1) are shown in purple, while those respecting only the FDR threshold are depicted in black and not significant genes (FDR > 0.1) in gray. Gene symbols highlight the top 30 genes (ranked by FDR). **(B)** Word cloud reporting for the same analysis DEGs gene symbol; word size is according to fold change magnitude, with genes with higher expression in 7Dup or in WBS in green and blue respectively. **(C)** STRING-based (https://string-db.org/) network reconstructed for the genes found modulated (FDR < 0.1, log2FC > 1) in at least one time-point by stage-wise differential expression analysis. In each visualization, the most relevant findings are highlighted among the DEGs identified in day 18, day 50 or day 100 COs. Node size and color represent the magnitude and direction of Log2FC at the specified stage, with genes more expressed in 7Dup and WBS in green and blue, respectively. Manual curation of functional annotation identified some functional relevant regions, highlighted in the network. Complete results are reported in supplementary figure S4A. **(D)** Volcano plot showing significantly up- and down-regulated proteins upon label-free quantification analysis, comparing expression levels in 7Dup and WBS samples, respectively (one CO/line, 3 lines/genotype in duplicate). Differential protein expression cutoff: FDR < 0.01 and Fold Change>1. **(E)** Barplots showing functional enrichment results for the Biological Process domain of gene ontology for up-regulated (p value < 0.05, FC>2) and down-regulated (p value < 0.05, FC < −2) proteins from 7Dup versus WBS comparison in Day 50 organoids. The top-8 categories according to p value are shown. **(F)** Venn diagram depicting the overlap between transcriptomics and proteomics results in 7Dup versus WBS day 50 cortical organoids. Features are selected as modulated with a conventional PValue < 0.05 and absolute log2FC > log2(1.25). **(G)** Scatterplot representing the relationship of the fold change in 7Dup vs WBS in the proteome (y axis) and in the transcriptome (x axis) for the subset of genes tested by both approaches. Features modulated in both conditions are highlighted in yellow and identified by their gene symbol; with the exception of *LYPLA2,* they are all modulated in the same direction at the transcript and protein level. DEGs found only in the proteome and in the transcriptome are visualized in purple and orange respectively. **(H)** Manually-curated functional annotation for subset of genes found modulated in the same direction at the transcript and protein level.

In order to identify the time-dependency of these transcriptional changes, we applied DEA separately at each CO differentiation stage and used the genes differentially expressed at least at one stage (FDR < 0.1 and Log2FC > 1 as absolute value) as input for the generation of stage-wise gene networks (**Figure S4A**). As expected, several genes from the 7q11.23 locus were consistently upregulated in 7Dup at all stages (**Figure 2C**). Other genes consistently upregulated in 7Dup in more than one stage were a group of poorly characterized zinc finger transcription factors (e.g., *ZNF229*, *ZNF300, ZNF560, ZNF578*). Focusing at each differentiation stage separately, the most outstanding observations included: i) at day 18, a modulation of cell adhesion and ECM components, mostly up-regulated in WBS COs. Of note, several of these genes (e.g., *COL12A*, *PCOLCE*, *BGN*, *NID2, EMILI1*) belong to a fetal cortex gene co-expression module characterized by non-monotonic expression pattern during cortex development and functionally associated to ECM (Cheroni et al., 2022); ii) at day 50, an up-regulation in WBS COs of histone protein genes as well as RG marker Nestin, consistent with the observed increase in proliferative cells in WBSs COs. This was accompanied at the same stage with the up-regulation in 7Dup COs of dorsal forebrain commitment drivers, including *FOXG1*, *LHX2*, *NEUROD6* and *TBR1*, whose trend of modulation was conserved in fold-change also at day 100 (**Figure S4A**); and, iii) the upregulation at day 100 of genes involved in synaptic transmission in WBS COs (e.g., *GRIK1*, *GRIA4*, *GRIN3A*). Of note, among the top upregulated genes in 7Dup COs at day 50 was *COL22A1*, a recently identified marker of a subset of layer III glutamatergic neurons, linked with human-specific evolutionary protracted developmental programs (Berg et al., 2021; Hodge et al., 2019).

In order to identify potential convergences between the transcriptional programs affected by 7q11.23 CNV and the causative genes of other neurodevelopmental disorders, we performed an overlap analysis of DEGs with public gene databases for neurodevelopmental syndromes (SFARI, Autism KB) and genetic diseases (OMIM, Orphanet and Decipher). This analysis retrieved significant overlaps for several gene sets, revealing a notable convergence between the genes transcriptionally modulated by 7q11.23 CNV and disease-causing genes, among which ASD and other neurodevelopmental disorders were prominently featured (**Figure S4B**).

Finally, reasoning that the neuronal imbalances observed at day 100 represented the output of earlier dysregulations during the mid-stage development, we focused proteome profiling on day 50, in order to uncover the upstream CNV-dependent protein expression changes that could account for later population imbalances. Analysis of differentially expressed proteins between 7Dup and WBS confirmed an up-regulation of proteins related to cell cycle and progenitor proliferation categories in WBS and, conversely, an up-regulation of neuronal maturation proteins in 7Dup (**Figure 2D,E**). Transcriptome/proteome overlap highlighted 30 common dysregulated genes/proteins, including a subset of the 7q11.23 members (higher in 7Dup) and a set involved in cell cycle up-regulated in WBS compared to 7Dup, further underscoring the symmetrically opposite prevalence of proliferative pathways in WBS and neurogenic programs in 7Dup.

### *GTF2I* gene dosage drives precocious neuronal differentiation in 7Dup COs

In order to elucidate the impact of progenitor imbalances on neuronal differentiation, we performed an in-depth inspection of the single cell data, focusing on the composition of the postmitotic neuron cluster (EN2), where we identified major differences between patient and control COs (**Figure 3A**). We used force-directed graph and Leiden clustering to identify the subpopulations composing the EN2 cluster (**Figure 3B**). This type of low dimensionality visualization tends to concentrate the most differentiated cells towards the external part of the graph. Benchmarking of COs single cell data against transcriptome from the human fetal cortex (Nowakowski et al., 2017) revealed the presence of transcriptional signatures of early neurons, Layer V-VI, Layer II-III, prefrontal, occipital and interneuron progenitors, highlighting the neuronal diversity that is recapitulated in COs (**Figure 3B**). A closer inspection of the distribution of cells in this neural compartment, at both day 50 and day 100, revealed marked differences between genotypes. Specifically, while cells from both WBS and CTL COs broadly overlap in areas with differentiated populations of excitatory and inhibitory neural lineages, 7Dup cells diverged further, showing increased abundance of postmitotic neurons (**Figure 3C**).

**Figure 3.**
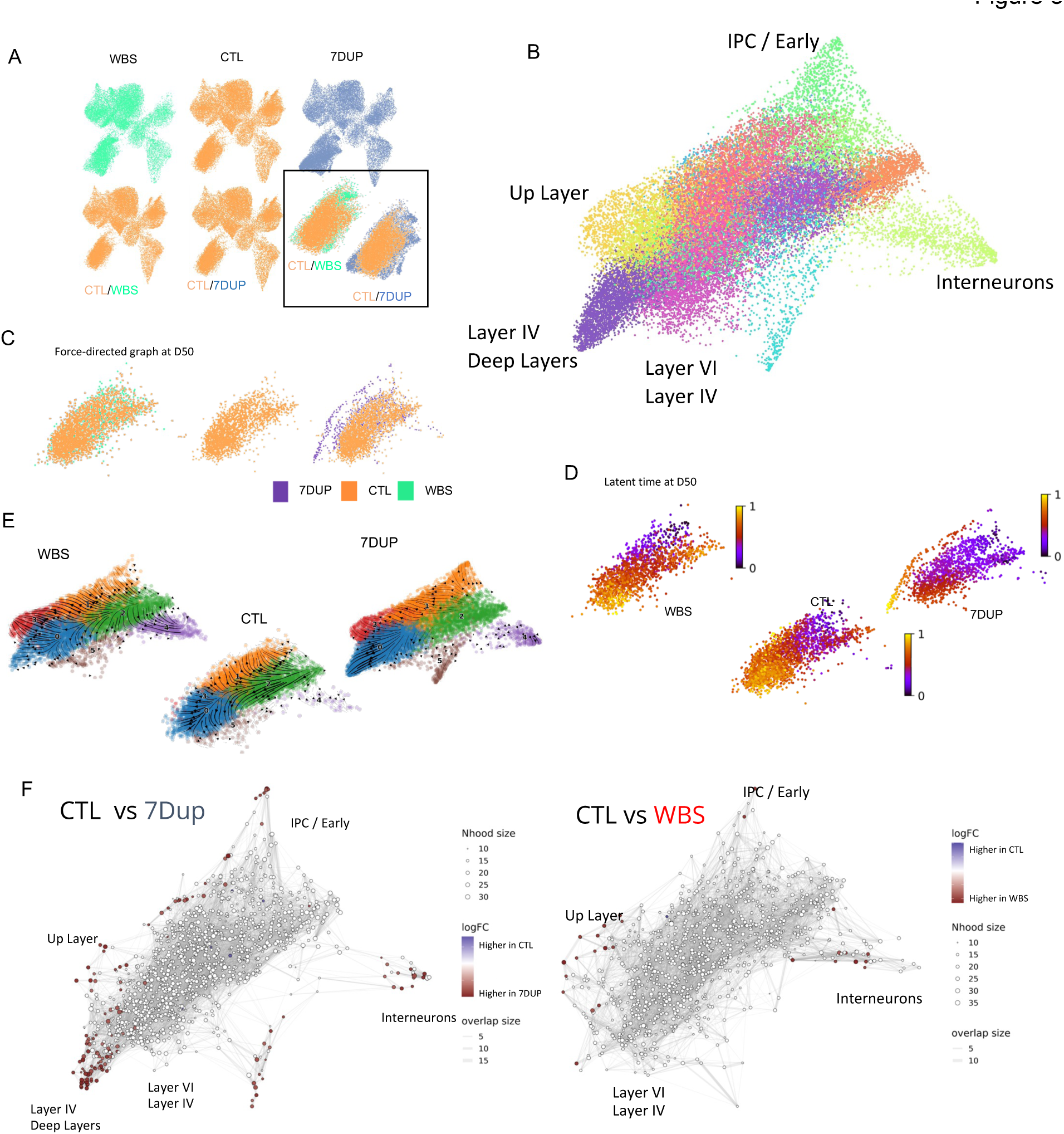
Gene dosage imbalances at 7q11.23 accelerate neuronal differentiation in 7Dup COs. **(A)** UMAP from (n = 97108) single cell transcriptomes from 21 biological samples, stratified for genotype (CTL, WBS and 7Dup from left to right, first line). UMAP in the second line shows the differences in cell distribution among genotypes, highlighting areas in the EN2 population that are unique to 7Dup and WBS, further highlighted in the zoom box (bottom right). **(B)** Force-directed graph drawing of only the EN2/IN populations (recalculating neighbors and distances sub selecting only EN2/IN populations), divided in subclusters and manually annotated. This dimensionality reduction highlights the most differentiated cell population (the pointed extremity of the graph, manually annotated). **(C)** Differences in cell distribution in force-directed graph for Day 50 organoids. WBS and CTL occupy the same space, while 7Dup has more mature cells. **(D)** Latent time calculated only on the EN2 and IN populations at Day 50. The starting cell is identified considering the total population. **(E)** Representation of the EN2 and IN populations developmental path in the 3 genotypes in force-directed graph, overimposed with RNA velocity. **(F)** Differential abundance of cells in different areas of the force-directed graph. The nodes are neighborhoods of cells aggregated based on RNA expression similarity, while graph edges depict the number of cells connected between adjacent neighborhoods. White nodes are neighborhoods that are not detected as differentially abundant (FDR > 1%), red dots, on the other end, are significant (FDR < 1%).

In order to inquire whether the increase of postmitotic neurons in 7Dup organoids (**Figure S3, 3C**) was the result of an accelerated neuronal differentiation program, we applied to the postmitotic neuron cluster the CellRank algorithm (Lange et al., 2022) that combines 3 sources of information (pseudotime, splicing and transcriptomic distance) to determine the probability of each neurogenic trajectory for each genotype and calculated the latent time, a pseudo-time representation that better defines the developmental trajectory. This analysis revealed a selective bias of 7Dup COs towards a neuronal cluster enriched in postmitotic neurons, which strikingly were present already at day 50 in 7Dup but absent in WBS and CTL (**Figure 3D**); this accelerated differentiation bias of 7Dup COs was further supported by the results of Velocyto algorithm (La Manno et al., 2018) in (**Figure 3E)**, where the lack of velocities in the center in 7Dup map indicates a more steady state of already differentiated cells while this is not the case in WBS and CTL. We then examined the differential abundance of cells in the force-directed graph representation (**Figure 3F**). The analysis uncovered a higher abundance of 7Dup cells in nodes representing mature neuronal populations. Collectively, the developmental trajectory analysis (**Figure 3D**), the velocyto analysis (**Figure 3E**) and the density of cells (**Figure 3F**) revealed that while cells from both WBS and 7Dup COs are enriched in clusters representing more differentiated neurons compared with CTL, including specific cortical layer (i.e., deeper and upper layer) neurons and interneurons, the exclusive presence of 7Dup COs cells in areas occupied by mature neurons (i.e., extremities of the graph) at day 50 indicates an accelerated neuronal differentiation in 7Dup (**Figure 3C-F**).

Given the pivotal role that *GTF2I* plays in the cognitive-behavioral manifestations in 7q11.23 CNV syndromes (Adamo et al., 2015; Crespi and Hurd, 2014; Gagliardi et al., 2003; Karmiloff-Smith et al., 2012) and its elevated expression in early and postmitotic neurons compared to its homologues GTF2RD1 and GTF2IRD2 (**Figure S6A-E**), we asked whether downregulating *GTF2I* in 7Dup COs would suffice to rescue the alterations in the developmental trajectories of the mature neuronal populations uncovered above. Indeed, COs generated from 7Dup iPSC lines constitutively expressing a *sh*RNA against GTF2I (7DupshGTF2I) (**Figure S6G,H**) showed reduced presence of mature neuronal populations, with a population composition resembling closely the one of control COs (**Figure 4A**). Likewise, RNA velocity confirmed that *GTF2I* downregulation in 7Dup COs leads to differentiation trajectories matching those of control COs (**Figure 4B**), suggesting that *GTF2I* dosage is key in the production of postmitotic neuronal populations in 7Dup individuals. We then once again applied CellRank to determine the probability of each neurogenic trajectory for each condition. First we identified all the possible endpoints for 7Dup, 7DupshGTF2I and control, the distribution of trajectory endpoints with a probability (P > 0.95, p-value < 0.05) revealed a robust bias to produce neurons enriched in Layer IV and deep layers markers (**Figure 4 C,D; S5 A,B**) in 7Dup COs at day 50, compared to controls. Remarkably, this bias was absent in COs with a constitutive knockdown of *GTF2I* (**Figure 4E**), thus confirming the central role of *GTF2I* in accelerated neuronal production in 7Dup COs.

**Figure 4.**
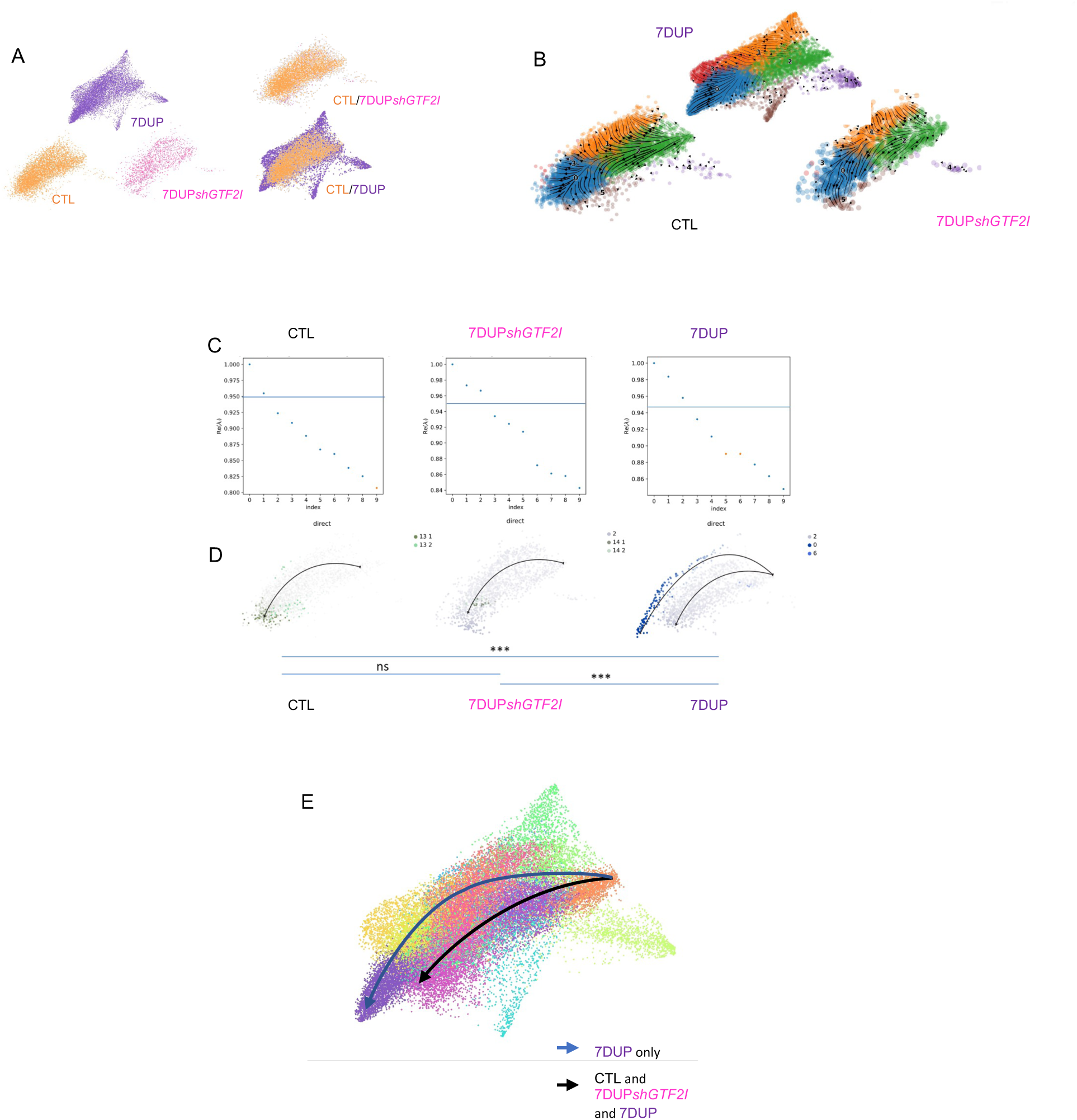
GTF2I duplication drives precocious neuronal differentiation in 7Dup COs. **(A)** Force-directed graph stratified for genotype (CTL, 7Dup and 7DupshGTF2I) from left to right, first line at D50. The second line shows the differences in cell distribution among genotypes, highlighting areas in the EN2 population that are unique of 7Dup and WBS, overimposed on the right. **(B)** Eigenvalue of the transition matrix stratified by genotype (CTL, 7Dup and 7DupshGTF2I). In the plot, each dot represents a potential differentiation trajectory, with y-axis quantifying the likelihood of solving the transition matrix (probability to be a proper differentiation trajectory); the blue line identifies the imposed probability threshold (0.95) applied to all the conditions. **(C)** Force-directed graph overimpose with RNA velocity, depicting the developmental path of different genotypes (CTL, 7Dup and 7DupshGTF2I). **(D)** Force-directed graph with color code as the probability of a cell to go to a certain endpoint, defined by the graph in panel C. Given the clustering used we have multiple clusters that identify the same populations. The significance is defined as the probability of going from the beginning, identified as the population with minimum of total pseudotime, to the endpoints in 7Dup. **(E)** Force-directed graph with a visual representation of the differentiation trajectory and populations that are involved in the process, only 7Dup passes in regions showing upper layer markers.

### Increased *Gtf2i* dosage drives accelerated neuronal differentiation and ASD-related behaviors in the mouse

Having defined *GTF2I* dosage as the critical effector of the altered neurodevelopmental trajectories uncovered in 7Dup, we asked whether this specificity could be leveraged *in vivo* to rescue 7Dup-associated cognitive-behavioral manifestations. To this end we resorted to murine models specifically recapitulating the *GTF2I* dosage of the 2 syndromes, i.e., carrying either 1 (Gtf2i^+/-^) or 3 (Gtf2i^+/Dup^) *Gtf2i* copies, alongside their wild type (WT) controls with the physiologic complement of 2 alleles (Mervis et al., 2012). First, we probed the extent to which these murine models featured equivalent alterations to those observed in human corticogenesis, in terms of *GTF2I* dosage-dependent alterations of the proliferative and mature neuronal compartments. Thus, we quantified the expression of proliferative and neuronal markers in the cortices of transgenic mouse embryos at E17.5. We found an increased expression of phospho-histone H3 (Phh3) in the Gtf2i^+/-^ embryos indicating a higher proportion of neuronal progenitors undergoing mitosis (**Figure 5A**), consistent with the findings from WBS COs, Ki67 was instead expressed more in Gtf2i^+/Dup^ embryos (**Figure 5B**), pointing to species-specific differences in the well-established distinct features of cell cycle phases captured by these two markers, with Phh3 expressed just prior to mitosis while Ki67 marking the entire cell cycle but was also expressed in non-actively proliferating cells (Colman et al., 2006; van Oijen et al., 1998). Consistently, also intermediate progenitor marker Tbr2 was significantly higher in WT compared with Gtf2i^+/Dup^ embryos (**Figure 5C**), whereas the increase in Gtf2i^+/-^ genotype *vis a vis* the Gtf2i^+/Dup^ did not reach significance. In terms of the postmitotic compartment, lower-and upper-cortical layer neuronal markers, Bcl11b and Cux1 respectively, were both robustly increased in Gtf2i^+/Dup^ embryos at E17.5 compared with Gtf2i^+/-^ and WT (**Figure 5D,E**).

**Figure 5.**
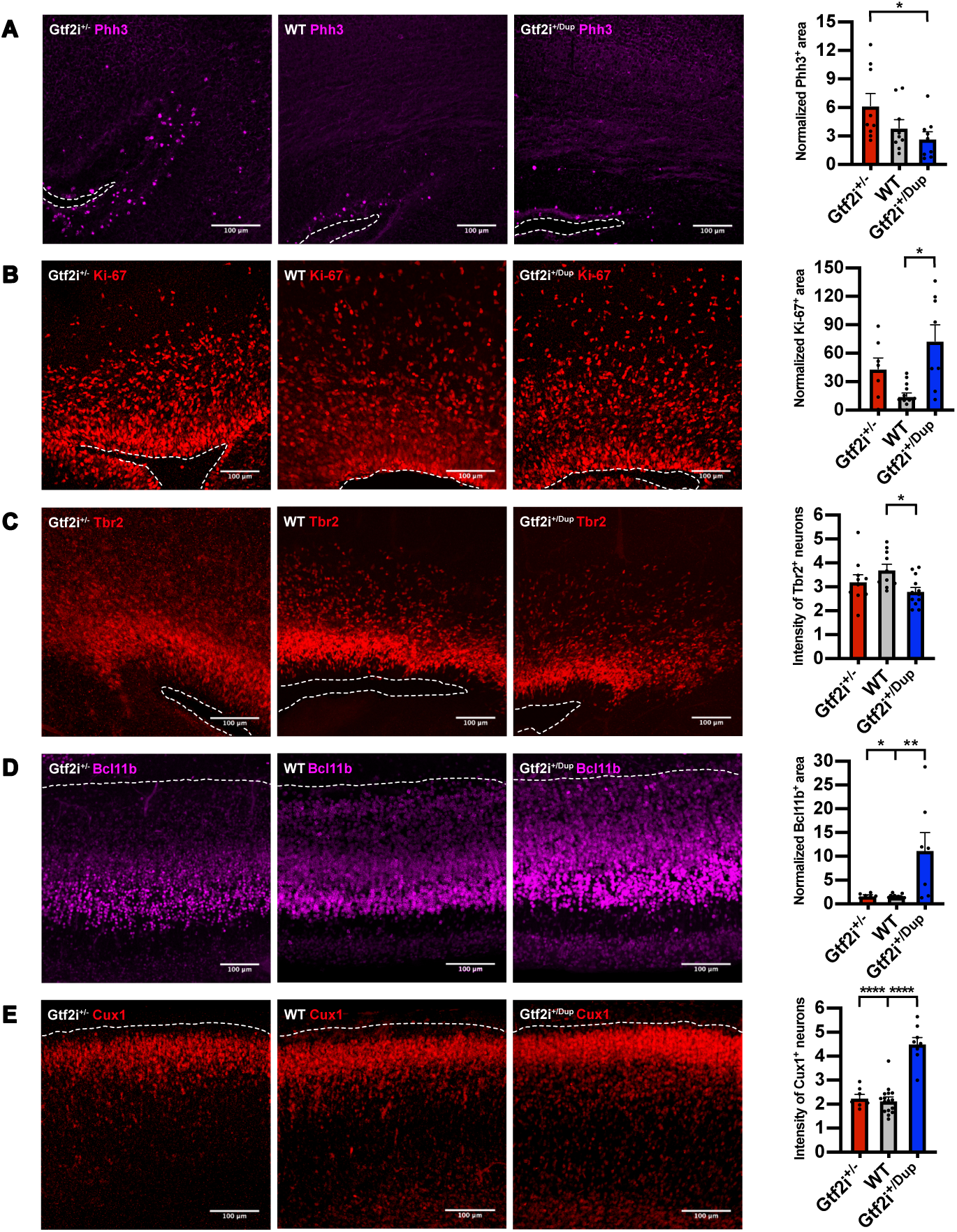
*Gtf2i* dosage drives accelerated neuronal differentiation in the mouse developing cortex. **(A)** (Left), Representative images of mitosis marker Phh3 expression in Gtf2i^+/-^, WT and Gtf2i^+/Dup^ E17.5 embryos. (Right), Quantification of Phh3 expression at E17.5 in the three genotypes. Gtf2i^+/-^ n=9 sections from 6 mice; WT n=9 sections from 5 mice; Gtf2i^+/Dup^ n=9 sections from 5 mice. **(B)** (Left), Representative images of cell cycle marker Ki-67 expression. (Right), Scatter plot with bar quantifying the Ki-67 expression at E17.5. Gtf2i^+/-^ n=7 sections from 4 mice; WT n=13 sections from 8 mice; Gtf2i^+/Dup^ n=8 sections from 4 mice. **(C)** (Left), Representative images of intermediate progenitor marker Tbr2. (Right), Scatter plot with bar quantifying Tbr2 expression at E17.5. Gtf2i^+/-^ n=11 sections from 6 mice; WT n=10 sections from 5 mice; Gtf2i^+/Dup^ n=12 sections from 5 mice. **(D)** (Left), Representative images of lower- layer cortical neuronal marker Bcl11b. (Right), Scatter plot with bar quantifying Bcl11b expression at E17.5. Gtf2i^+/-^ n=7 sections from 4 mice; WT n=16 sections from 8 mice; Gtf2i^+/Dup^ n=8 sections from 4 mice. **(E)** (Left), Representative images of upper-layer cortical neuronal markers Cux1. Right, Scatter plot with bar quantifying Cux1 expression at E17.5. Gtf2i^+/-^ n=7 sections from 4 mice; WT n=16 sections from 8 mice; Gtf2i^+/Dup^ n=8 sections from 4 mice. All data are shown as mean±SEM. Statistical analyses were performed using one-way ANOVA followed by Tukey’s multiple comparisons test. Significance level was set to p<0.05. *p<0.05; **p<0.01; ***p<0.001; ****p<0.0001.

To further probe the *Gtf2i* dosage-sensitive corticogenesis phenotypes, we electroporated *in utero* pCAG-GFP at E14.5 and analyzed cortices at E17.5 or P7.5 (**Figure S7A**). Neurons from Gtf2i^+/Dup^ mice at E17.5 had an aberrant orientation and were stalled between SVZ and IZ, failing to migrate into the cortical plate (**Figure S7B,C**). We also examined the neuronal morphology postnatally at P7.5 mice (**Figure S7D**). Both mutants had an altered neuronal morphology compared with WT, with fewer dendritic branches and lower number of dendrites (**Figure S7E-G**). Together, these results validate *in vivo* the robust impact of an increased *GTF2I* dosage on the neuronal proliferation/differentiation dynamics during corticogenesis, while uncovering additional alterations in migration and branching (the latter shared with the hemideleted dosage), underscoring how even at the level of a single gene the 7q11.23 CNV exerts composite effects spanning the range of shared versus symmetrically opposite endophenotypes.

Next, having elucidated the impact of *GTF2I* dosage on human corticogenesis and validated partially equivalent phenotypes in murine models recapitulating its CNV, we set out to define its contribution to ASD-relevant behavioral phenotypes employing the three-chamber sociability test (Moy et al., 2004; **Figure 6A,B**). In the first part of the test, we quantified the time that male or female *Gtf2i*^+/Dup^ and WT littermates spent with an object or a conspecific. In contrast to WT that spent more time with the conspecific, *Gtf2i*^+/Dup^ showed either no preference for the object versus the conspecific or spent more time with the object (for male and female mice, respectively, **Figure 6C,D; Figure S8A,B**). During the second part of the test, mice had a choice between spending time with a familiar or a novel conspecific. As expected, WT mice spent significantly more time with a novel conspecific, whereas *Gtf2i*^+/Dup^ showed no preference for the familiar versus the novel mouse (**Figure 6E,F; Figure S8C,D**). The robustness and significance of these phenotypes, particularly salient given the outbred nature of the *GTF2I* strains (CD-1 background), provided the rationale for testing rescue strategies, given the precedents of other syndromes such as Fragile X, in which the pharmacological rescue of behavioral phenotypes in adult animals had redefined the horizon of treatment options for neurodevelopmental disorders (Licznerski et al., 2020; Michalon et al., 2012). We had previously established that *GTF2I* recruits a chromatin repressive complex featuring histone H3 lysine 4 demethylase LSD1 and that LSD1 inhibition relieves aberrant gene repression brought about by increased *GTF2I* dosages in differentiating 7Dup iPSC (Adamo et al., 2015). We thus reasoned that LSD1 inhibition *in vivo* could be a viable strategy to rescue ASD-like *GTF2I*-dependent behavioral phenotypes and targeted LSD1 with a specific inhibitor (Vianello et al., 2016) that readily crosses the blood brain barrier (Faletti et al., 2021) in *Gtf2i*^+/Dup^ male mice, given the higher prevalence of ASD in males (Loomes et al., 2017). As shown in (**Figure 6G-J**), administration of LSD1 inhibitor via oral gavage for 2 weeks, rescued deficits in social preference and social novelty in *Gtf2i*^+/Dup^ mice. Blood samples taken after LSD1 inhibitor treatment confirmed that the treatment did not cause hematopoietic toxicity (Serfilippi et al., 2003) (**Figure S9A-C**). Next, we sought to test an interval treatment regimen that would have even greater translational relevance, especially for the prospect of chronic administration starting from a young age. Though we used an irreversible LSD1 inhibitor, it was in fact not known whether and for how long the beneficial impact on behavior would last. We thus devised a regimen in which, following LSD1 inhibitor treatment, we allowed mice 2 weeks of rest to remove the LSD1 inhibitor from the system and then tested them again in the three-chamber sociability apparatus. Remarkably, the beneficial impact of LSD1 inhibition on ASD-relevant behaviors was evident even after the 2 weeks “washout” period without detectable hematopoietic toxicity (**Figure. 6K-N; Figure S9D-F**). Collectively, these results show that increased *Gtf2i* gene dosage underlies ASD-like behaviors and that its interference by LSD1 inhibition is a viable path for therapeutic intervention in 7Dup.

**Figure 6.**
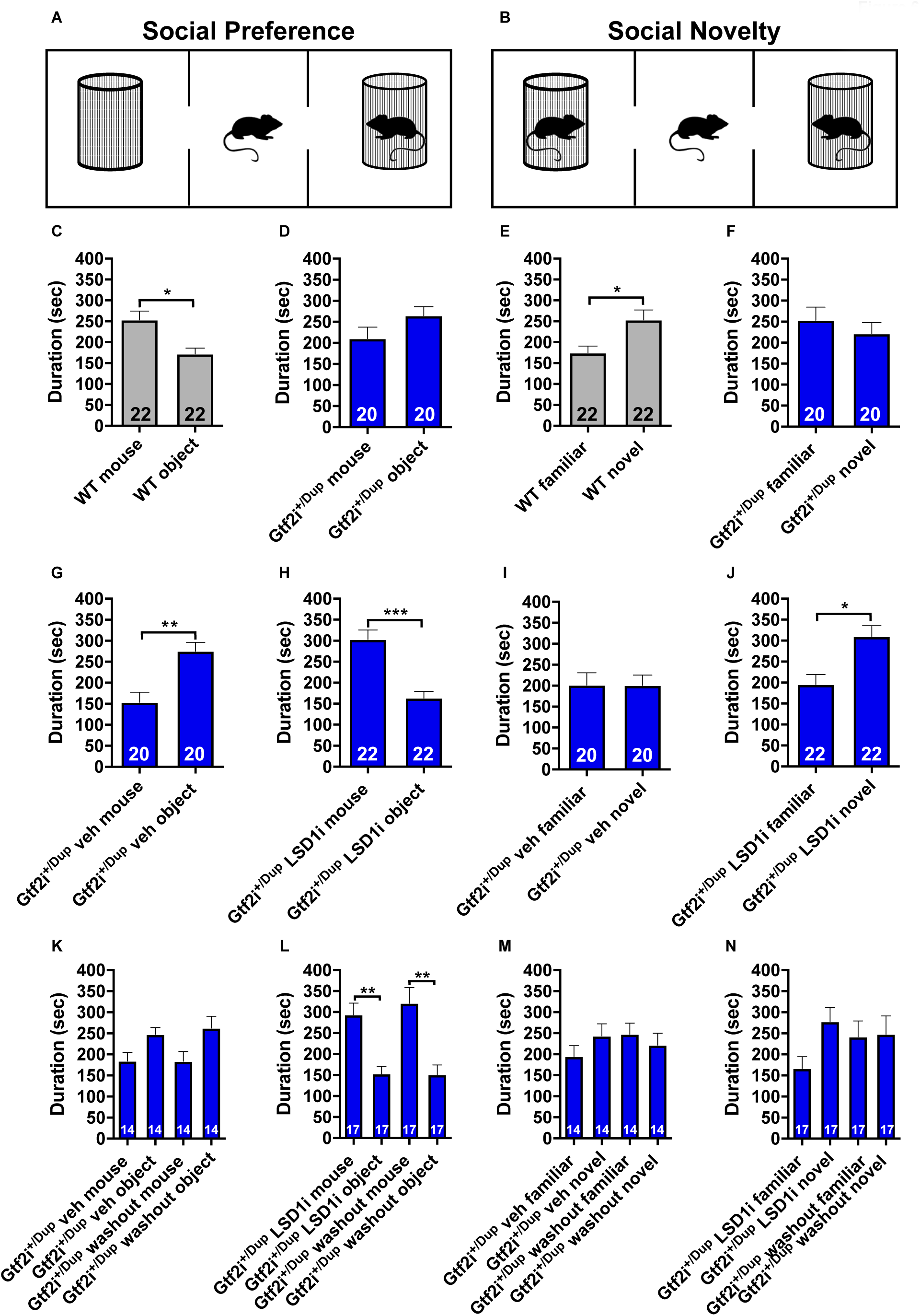
Inhibition of LSD1 rescues ASD-like phenotypes in Gtf2i^+/Dup^. **(A,B)** Schematic representation of the three-chamber sociability apparatus for measuring social preference and social novelty, respectively. **(C-F)** Bar plots with dots depicting the time spent with a conspecific vs object and with a novel vs familiar mouse at baseline, in Wt (n=22) and Gtf2i^+/Dup^ mice (n=20). **(G-J)** Bar plots with dots depicting the time spent with a conspecific vs object and with a novel vs familiar mouse in Gtf2i^+/Dup^ mice following 4 oral gavage administrations (2 times/week over 2 weeks) of vehicle (n=20) or LSD1 inhibitor (10mg/kg; n=22). **(K-N)** Bar plots with dots depicting social preference and social novelty results in Gtf2i^+/Dup^ mice tested after the 4^th^ administration of vehicle (n=14) or LSD1 inhibitor (n=17) and after a 2 week “washout” period. Data shown as mean±SEM. Statistical analyses were performed using paired Student’s t-test followed by Holm-Bonferroni correction for multiple testing. Significance level p<0.05. *p<0.05; **p<0.01; ***p<0.001.

### LSD1 inhibition affects neurodevelopmental and synaptic organization molecular pathways in the cortex of Gtf2i^+/Dup^ mice

Finally, to define the molecular underpinnings of the efficacy of LSD1 inhibitor, we treated Gtf2i^+/Dup^ mice with either 1 or 4 doses of LSD1 inhibitor and collected the cortices i) 24 hours after for the single dose (acute), or ii) 2 hours following the 4^th^ dose and immediately after the behavioral test (chronic; **Figure 7A**). We focused our analysis on prefrontal and somatosensory cortices, two brain regions implicated in ASD pathophysiology (Parikshak et al., 2013; Willsey et al., 2013). As expected, inhibition of the transcriptional repressor LSD1 lead to an overall up-regulation of genes. In particular, acute treatment resulted in 114 DEGs (FDR<0.1, 99 up-regulated and 15 down-regulated) (**Figure 7B,C**). Prominent up-regulated genes are implicated in neurodevelopment and neurodevelopmental syndromes, including *Ahi1*, which is required for cerebellar and cortical development and causes Joubert syndrome (Dixon-Salazar et al., 2004), *Irf2bpl,* which is associated with neurodevelopmental disorder with regression and plays a role in development of CNS via negative regulation of Wnt signaling (Tran Mau-Them et al., 2019), *Doc2a,* implicated in schizophrenia and ASD (Glessner et al., 2010), and *Arvcf* linked to DiGeorge syndrome and schizophrenia (Sanders et al., 2005). On the other hand, chronic LSD1 inhibition uncovered 54 DEGs (FDR<0.1), 46 of which up-regulated (**Figure 7D,E**). Among the dysregulated genes was a consistent upregulation of protocadherin gene family members (i.e., *Pcdhgb5, Pcdhb6, Pcdhb7, Pcdhb11, Pcdhga5*), as in 7Dup COs. Protocadherins are calcium-dependent cell adhesion proteins, which establish and maintain synapse formation and neuronal connections (Weiner and Jontes, 2013) and their dysregulation has been strongly implicated in neurodevelopmental disorders (Flaherty and Maniatis, 2020). These findings suggest that inhibition of Gtf2i-LSD1 complex causes a two-tiered effect on the mouse’s cortex. In the short-term triggers an upregulation of genes critical for neurodevelopment. Subsequently, sustained inhibition of the complex initiates remodeling of synaptic organization by increasing the expression of protocadherins, which in turn are critical for sociability (Flaherty and Maniatis, 2020).

**Figure 7.**
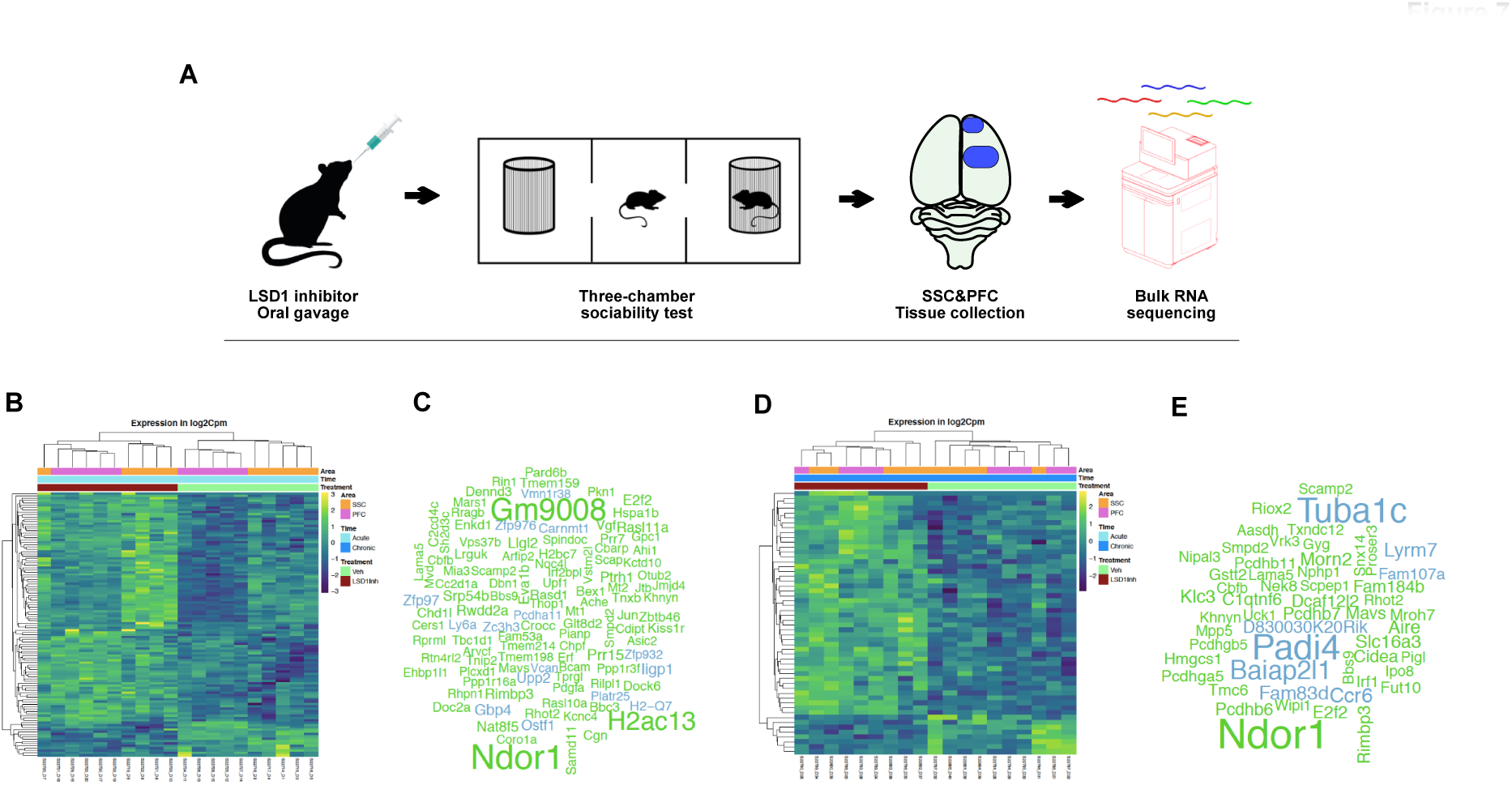
LSD1 inhibition affects neurodevelopment and synaptic organization in the cortex of Gtf2i^+/Dup^ mice. **(A)** Schematic representation of the experimental flow. Gtf2i^+/Dup^ mice received either a single (acute) or 2 administrations of LSD1 inhibitor/week over 2 weeks (semi-chronic) followed by three-chambered sociability test. Prefrontal (PFC) and somatosensory (SSC) cortices were dissected, flash frozen and RNA extracted for bulk transcriptomic profiling. **(B-C)** Heatmap and word cloud respectively depicting the expression levels (as z-scores) and gene symbol (green: up-regulated; blue: down-regulated) of the 114 genes identified as DEGs following a single administration of LSD1 inhibitor compared to vehicle in pre-frontal and somatosensory cortex. Vehicle, n=5 mice and LSD1 inhibitor, n=5 for both cortical areas. **(D-E)** Chronic administration of LSD1 inhibitor results in 54 DEGs as shown in the heatmap and the word cloud. Genes from the protocadherins family were prominently upregulated as a result of chronic LSD1 inhibition. Note in both acute and chronic paradigms, the upregulation of most genes after inhibition of the transcriptional suppressor LSD1. Vehicle, n=5 mice for both areas; LSD1 inhibitor, n=5 mice for SSC and n=4 mice for PFC.

## Discussion

WBS and 7Dup syndromes are paradigmatic of the subset of neurodevelopmental disorders whose symmetrically opposite CNV are mirrored in symmetrically opposite phenotypes, thereby affording a unique glimpse into the dosage-sensitive mechanisms that underlie foundational aspects of the human condition such as sociability and language (López-Tobón et al., 2020). This uniquely explanatory potential has remained however largely unfulfilled due to three main limitations. First, while genetic evidence provides the unequivocal causal links between gene dosage and high-order human phenotypes, the neurodevelopmental antecedents that mediate those links are still mostly unscrutinized. Hence the neuroconstructivist framework that emphasized developmental constraints and the need to resolve both typical and atypical development in terms of the underlying trajectories, and that owes so much of its articulation precisely to WBS, is yet to be translated in molecular terms (Karmiloff-Smith et al., 2018). Second, tracing the neurodevelopmental antecedents mediating those causal links requires the combination of multiple experimental systems, so as to bridge the distance between higher-order functions and the underlying gene dosage. Yet previous studies employed, separately, either animal models or 2D *in vitro* cellular models profiled in bulk, thus precluding both the tracing of neurodevelopmental trajectories at high resolution and the cross-validation between experimental systems needed to bridge across endophenotypes unfolding at different layers of biological function. Third, the cis-epistasis of the 7q11.23 CNV, in which the simultaneous dosage imbalances of several genes within the CNV can potentially contribute to phenotypes, has made its modelling notoriously challenging. Thus, despite recent successes by us and others in delineating the roles of 7q11.23 locus’ genes in phenotypes relevant for disease pathophysiology, including, *FZD9* (Adamo et al., 2015; Barak et al., 2019; Chailangkarn et al., 2016; Tebbenkamp et al., 2018; Zanella et al., 2019), the impact of 7q11.23 gene dosage alterations specifically on neurodevelopment and behavioral manifestation remain unknown. Here, we combine evidence from COs and transgenic mice to identify *GTF2I* and its axis as a central hub of neuronal maturation that underlies complex behavioral manifestations.

Analysis of single cell transcriptome of neural progenitors and postmitotic neurons revealed a set of genotype-specific asymmetries, including an early production of mature neurons in 7Dup COs, whereas the developmental trajectories revealed divergent paths in the two genotypes, favoring upper-layer neurons in 7Dup and lower-layer neurons in WBS, while MGE-derived interneuron progenitors were similar in both genotypes. Definitions of differentiation trajectories through RNA velocity, which accurately predicts the future state of cells, revealed that these alterations in cortical neuronal fates are the result of heterochronic neuronal differentiation (i.e. precocious in 7Dup, delayed in WBS), which is a first symmetrically opposite endophenotype distinguishing, as neurodevelopmental antecedent, the divergent behavioral manifestations of WBS and 7Dup. Critically, we found consistent symmetrically opposite neuronal differentiation dynamics in WBS and 7Dup using 2D neuronal cultures in the accompanying article in the same issue (Mihailovich et al.), corroborating the timing of neuronal differentiation-as a critical yet experimentally tractable phenotype-across experimental models and systems. Asynchronous generation of specific neuronal populations is an emerging cellular mechanism underlying the convergent impact of autism genes on tractable neuronal phenotypes, as we and others have recently shown (Mariani et al., 2015; Paulsen et al., 2022; Villa et al., 2022). Here, we reveal that this cellular mechanism extends beyond high confidence autism risk genes to symmetrically opposite CNV, thereby providing a high-resolution cellular framework to annotate the neurodevelopmental trajectories of the sociability spectrum (López-Tobón et al., 2020). This heterochronic neuronal differentiation was largely corroborated by our findings in mice, where we found an increased number of postmitotic neurons in Gtf2i^+/Dup^ compared with Gtf2i^+/-^ mice.

*GTF2I* preferentially binds promoters and drives both basal and signal-induced transcription (Roy, 2001), and moreover it targets and interacts with critical genes for embryonic development and patterning, cell growth and proliferation, including *SMAD2*, *ERK* and *JAK2* (Kim and Cochran, 2000, 2001; Ku et al., 2005). The formation of transcriptional suppressive complex with the well-established pluripotency regulator LSD1 (Adamo et al., 2011), as we previously showed (Adamo et al., 2015), lent further support to the notion that its effects most likely involve transcriptional programs critical for fate specification and neuronal differentiation. Our findings now show that epigenetic drugs targeting transcriptional regulation can be a viable strategy for neurodevelopmental disorders that have been so far notoriously difficult to treat. LSD1 inhibitors are under intense investigation for their promising efficacy in various human malignancies (Maiques-Diaz and Somervaille, 2016), and more recently LSD1 inhibitors, albeit different from the one we used here, proved efficacious in rescuing ASD-like behaviors in mouse models with deficient *Shank3* and *Cul3* (Rapanelli et al., 2022). The current data, as well as our recent report on the efficacy of HDAC inhibitors regulating *GTF2I* expression (Cavallo et al., 2020), highlight the potential of drug repurposing for neurodevelopmental disorders with extremely limited treatment options.

Interestingly, while the knockdown of *GTF2I* rescues the accelerated production of excitatory neurons in 7Dup, it had only a negligible impact on the asymmetry of progenitor production, underscoring how, in the context of a complex cis-epistatic CNV, the longitudinal dissection of endophenotypes serves to define single gene contributions that can be highly specific and time-dependent. This also paves the way to further studies aimed at dissecting how other genes from the same locus, acting alone or in combination with *GTF2I*, bring about distinct aspects of the endophenotypes we have now charted.

Animal and population studies have attributed a critical role to *GTF2I* in sociability phenotypes (Jabbi et al., 2015; Li et al., 2009; López-Tobón et al., 2020; Procyshyn et al., 2017; vonHoldt et al., 2017). Protocadherins on the other hand, are synaptic cell adhesion molecules (sCAMs) that play critical roles in synapse organization and function (Bourgeron, 2015; Redies et al., 2012). Not surprisingly, mutations in protocadherins lead to aberrant neuronal communication and function, which in turn affect higher cognitive processes, including sociability (Taylor et al., 2020). Bulk RNAseq results revealed a modulation of protocadherins in 7Dup COs and Gtf2i^+/Dup^ mice following LSD1 inhibition, suggesting a convergence across species and experimental models on a potential *GTF2I* dosage-protocadherins molecular axis mediating sociability.

Overall, these data provide an unprecedented molecular dissection of the 7q11.23 gene dosage alterations impact on neurodevelopment and behavioral phenotypes and uncover the GTF2I-LSD1 axis as a central mediator of 7q11.23 disease pathophysiology that is amenable to therapeutic intervention.

## Methods

### iPSC culture

We used 13 control and patient-derived iPSC lines that we generated previously and reported in (Adamo et al., 2015; Cavallo et al., 2020) (**see also supplementary table 2**). Relevant ethics approvals are referred to in the original publications reporting their first use and/or derivation. iPSC lines were cultured on plates coated with human-qualified Matrigel (Corning; dilution 1:40 in DMEM-F12) in mTeSR (StemCell Technologies) or TeSR-E8 medium (Stemcell Technologies). Cells were passaged with ReLeSR (StemCell Technologies) or with Accutase (Sigma). For single-cell culture, cells were resuspended in mTeSR or TeSR-E8 medium supplemented with 5 μM Y-27632 (Rock inhibitor (RI), Sigma).

### Generation of cortical organoids

Cortical brain organoids were generated by following the modified protocol described in (López-Tobón et al., 2019; Paşca et al., 2015). Since we optimized the protocol to avoid the use of mouse embryonic fibroblast (MEF) for stem cell culture, the following procedures were followed; when the hiPSC line reached 80% confluency in a 10 cm dish, colonies were dissociated with Accutase and centrifuged. After resuspension in TeSR/E8 medium supplemented with 5uM ROCK inhibitor (Corning) cells were counted with a TC20 automated cell counter (Biorad) and resuspended to get a final concentration of 2 x 105 cells/mL. 100 μl/well of cell suspension were seeded into ultra-low attachment PrimeSurface 96-well plates (SystemBio) and the plates were centrifuged at 850 rpm for 3 minutes to promote the formation of embryoid bodies (EB). The day of the EB generation is referred to as day −2. On day 0 neural induction media was added, consisting of of 80% DMEM/F12 medium (1:1), 20% Knockout serum (Gibco), 1 mM non-essential amino acids (Sigma), 0.1 mM cell culture grade 2-mercaptoethanol solution (Gibco), GlutaMax (Gibco, 1:100), penicillin 100 U/ml, streptomycin (100 μg/ml), 7 μM Dorsomorphin (Sigma) and 10 μM TGFβ inhibitor SB431542 (MedChem express) for promoting the induction of neuroectoderm. From day 0 to day 4, medium change was performed every day, while on day 5 neuronal differentiation medium was added, consisting of neurobasal medium (Gibco) supplemented with B-27 supplement without vitamin A (Gibco, 1:50), GlutaMax (1:100), penicillin 100 U/ml, streptomycin (100 μg/ml), 20 ng/ml FGF2 (Thermo) and 20 ng/ml EGF (Thermo) until day 25. From day 25-43 the neuronal differentiation medium was supplemented with 20 ng/ml BDNF and NT3 (Peprotech). From day 43 onwards no growth or neuronal maturation factors were added to the medium.

### Western Blot

Organoids were snap-frozen in dry ice and lysed in SDS buffer [100mM Tris-HCl (pH 7.6), 20% glycerol and 4.7% SDS] denatured at 95C° for 10 minutes and sonicated for 15 seconds. Protein extracts (30 to 50 ug) were mixed with Laemmli loading buffer, denatured for 5 minutes 95C°, electrophoresed in 10% SDS-PAGE gel and transferred to PVDF membranes. The membranes were blocked in TBST [50 mM tris (pH 7.5), 150 mM NaCl, and 0.1% Tween 20] and 5% milk for 1 hour and incubated overnight with GTF2I (BD Biosciences) and GAPDH (Millipore) primary antibodies (see supplementary table 1). Blots were developed with the ECL Prime Western Blotting Detection Reagents (Sigma-Aldrich) and bands imaged and quantified using the ChemiDoc system (Bio-Rad).

### Bulk Transcriptome Analysis

At the designated time-point, cortical organoids were washed once with PBS and snap-frozen in dry ice. Total RNA was isolated with the RNeasy Micro Kit (QIAGEN, Hilden, Germany) according to the manufacturer’s instructions. RNA was quantified with Nanodrop and then the integrity was evaluated with Agilent 2100 Bioanalyzer. A TruSeq Stranded Total RNA LT Sample Prep Kit (Illumina) was used for library preparation starting from 500 ng of total RNA for each sample. Sequencing was performed with the Illumina NOVAseq 6000 platform, with an average depth of 35 million 50 bp paired-end reads per sample.

#### Cortical Organoids

RNAseq FASTQ data were quantified at the gene level using Salmon (version 0.8.2). GRCh38 Genecode 27 was used as reference for quantification and annotation. Protein-coding genes were selected for downstream investigations.

Differential expression analyses (DEA) comparing 7Dup versus WBS cortical organoids were performed by edgeR either on the complete organoid cohort (33 samples: 3 WBS, 4 CTL and 3 7Dup for Day18 organoids; 4 WBS, 4 CTL and 4 7Dup for Day50 organoids; 3 WBS, 5 CTL and 3 7Dup for Day100 organoids) or selecting by differentiation stage. For the complete cohort analysis, genes with expression levels higher than 2 cpm in at least 4 samples were tested for differential expression; the information about organoid differentiation stage was used as a covariate in the statistical model. DEGs were selected imposing a threshold of FDR < 0.1 and absolute log2FC > 1.

Similarly, stage-wise DEAs were performed comparing 7Dup versus WBS and selecting genes with an expression level higher than 2 cpm in at least 3 samples for Day18 and Day100, and in at least 4 samples for Day 50. 315 genes that were found significantly modulated (FDR < 0.1, absolute log2FC > 1) in at least one time-point were selected for network reconstruction by retrieving STRING database connections (https://string-db.org/) and generating the network in Cytoscape (Shannon et al., 2003). Node shape highlights the stage in which the genes are significantly modulated: diamond for Day18, circle for Day50, square for Day100 and hexagon for genes significant in more than one stage. Identification of functionally-relevant regions in the network was performed by manual curation of node biological function.

The following knowledge bases were considered for the overlap analysis: MIM (Mendelian Inheritance in Man), Orphanet, Decipher, SFARI Genes, and AutismKB. MIM, Orphanet and Decipher genes were obtained by interrogating version 100 of Biomart (www.ensembl.org/biomart/martview/). SFARI genes and AutismKB were downloaded respectively from https://gene.sfari.org/database/human-gene/ and http://db.cbi.pku.edu.cn/autismkb_v2/download.php in April 2020. Overlap between these gene sets and differentially expressed genes at day18, day50 or day100 (either considered together or divided in up-regulated and down-regulated) was tested by GeneOverlap R library (version 1.14.0). Overlaps were considered significant with an odds ratio higher than 1 and p-value lower than 0.1.

Where not otherwise specified, analyses were performed in R, version 3.4.4

### LC–MS/MS analysis and raw data processing

In all cases, sample preparation was performed according to the in-StageTip protocol (Kulak et al., 2014). Briefly, samples (one CO/cell line, 3 cell lines/genotype in duplicate) were incubated in PreOmics Lysis Buffer (catalogue number P.O. 00001, PreOmics) for cysteine reduction and alkylation, followed by a protein denaturation step at 95 °C for 10 min. Then, proteins from each sample were digested by adding the Trypsin/LysC mix (Digestion buffer, PreOmics) at 1:50 enzyme-to-protein ratio, at 37 °C overnight. Proteolytic peptides were eluted from the solid-phase extraction material and dried out completely in a SpeedVac centrifuge (Eppendorf). Finally, peptides were re-suspended in 5 µL of Load Buffer (PreOmics) and analysed by nano- RPLC-MS/MS using an EASY-nLC 1200 (Thermo Fisher Scientific, cat. no LC140) connected to a Q-Exactive HF instrument (Thermo Fisher Scientific) through a nano-electrospray ion source. In all cases, the nano-LC system was operated in one-column setup with an EasySpray PEPMAP RSLC C18 column (Thermo Fisher Scientific) kept at the constant temperature of 45°C. Solvent A was 0.1% formic acid (FA) and solvent B was 0.1% FA in 80% Acetonitrile (ACN). Peptides were separated along a 64-min gradient of 3–30% solvent B, followed by a gradient of 30–60% for 10 min and 60–95% over 5 min, at a flow rate of 250 nL/min. The Q-Exactive was operated in the data-dependent acquisition (DDA) mode, to automatically switch between MS and MSMS mode. The 15 most intense peptide ions with charge states ≥2 were sequentially isolated to a target value of 3e6 (Top15). Spray voltage was set to 1.7 kV, s-lens RF level at 50, and heated capillary temperature at 275 °C. Selected target ions were dynamically excluded for 20 seconds and all experiments were acquired using positive polarity mode. The MS spectra (from m/z 375-1550) were analysed in the Orbitrap detector with resolution R=60,000 at m/z 200. MS2 data was acquired at R=15,000 resolution and an ion target value of 1e5. Higher-energy collisional dissociation (HCD) fragment scans was acquired using 1.4 m/z isolation width and normalized collision energy of 28. The maximum allowed ion accumulation times were 20ms for full scans and 80ms for MSMS.

The acquired raw MS data were analysed using the integrated MaxQuant version 1.6.2.3 (Tyanova et al., 2016a), using the Andromeda search engine (Tyanova et al., 2016b) and the Human Fasta Database downloaded from UniprotKB (74470 Entries) was used. Carbamidomethylation of Cysteine was set as a fixed modification. Enzyme specificity was set as carboxy-terminal to arginine and lysine as expected, using Trypsin and LysC as proteases. A maximum of two missed cleavages was allowed. Peptide identification was carried out with the Andromeda algorithm with an initial precursor mass deviation of up to 7 ppm and a fragment mass deviation of 20 ppm. False discovery rate (FDR) for both peptide and proteins was set to a maximum of 1% for identification. All proteins and peptides matching the reversed database were filtered out. The LFQ intensity calculation was enabled, as well as the match between runs (MBR) feature (Cox et al., 2014). The “protein groups” output file from MaxQuant was analyzed using Perseus software (Tyanova et al., 2016b). Briefly, no imputation was used, and data were filtered in order to have at least 3 valid values in at least one group. To identify significant regulated proteins an FDR=0.05 was imposed in the t-test analysis.

### Transcriptome-Proteome comparison

Results from the proteomics profiling were compared with the transcriptional changes identified at Day50 (7Dup vs WBS DEA). The comparison between the two approaches was performed by selecting features (genes or proteins) measured by both techniques and discarding those with duplicated gene symbols. Significant modulation has been defined according to the following thresholds: PValue < 0.05 and FC > 1.25 (absolute value); overlap was tested with a Fisher test and visualized by Venn diagram (R package VennDiagram version 1.6.20, https://cran.r-project.org/package=VennDiagram) and by custom scatter plot.

Proteomic functional analysis: Gene ontology enrichment analysis for the Biological Process domain was performed on the proteins identified as changed in 7Dup versus WBS with a conventional PValue < 0.05 and an absolute log2FC value of at least 1. Analysis was performed by TopGO (version 2.30.1,(Alexa et al., 2006), relying on Fisher test and Weight01 method to take into account ontology hierarchy; node size was set at 15. Cut offs of 0.01 on the p-value and 1.75 on the enrichment threshold were applied to select significantly enriched GO terms. The top-8 categories according to PValue for up-regulated and down-regulated genes were visualized by barplots.

### Overlap analysis with gene-disorder knowledge bases

MIM (gene-disorder association for genetic diseases), Orphanet (gene-disorder association for rare diseases) and Decipher knowledge bases were retrieved from Mart (Biomart version 100, www.ensembl.org/biomart/martview/). The Simons Foundation Autism Research Initiative (SFARI) gene list was downloaded from the SFARI website (https://gene.sfari.org/database/human-gene/, downloaded in April 2020). The evidence-based knowledge base for autism AutismKB was retrieved from (http://db.cbi.pku.edu.cn/autismkb_v2/download.php, downloaded in April 2020). Overlap analysis was performed between each considered knowledge base and the results of stage-wise DEA (7Dup vs WBS comparison), selecting DEGs with FDR < 0.1 and FC > 2 and testing them either as a single set or split in up and down-regulated genes. Overlap analysis was performed using GeneOverlap R package (version 1.14.0, http://shenlab-sinai.github.io/shenlab-sinai/) and considering the tested genes for each DEA as universe. Results were visualized by dotplot, reporting the number of overlapping genes for Odds Ratio (OR) > 1 and the dot for pvalues < 0.1. Dot size and color represent the p-value and the OR respectively.

### Single cell transcriptome analysis

Organoids collected from different genotypes and time points were dissociated by incubation with a solution of Papain (30U/mL, Worthington LS03126) and DNAseI (3U/uL, Zymo Research) for 30–45 min depending on organoid size. Dissociated suspensions were passed once through 0.4 mm Flowmi™ cell strainers, resuspended in PBS and counted using TC20 automatic cell counter (Biorad). Resulting single-cell suspension was mixed with RT-PCR master mix at a density of 1000 cells/μl and loaded together with Chromium Single-Cell 3′ gel beads and partitioning oil into a Chromium Single Cell 3′ Chip. The gel beads were coated with unique primers bearing 10x cell barcodes, unique molecular identifiers (UMI) and poly(dT) sequences. The chip was then loaded onto a Chromium instrument (10x Genomics) for single-cell GEM generation and barcoding. Amplified cDNAs were fragmented, and adapter and sample indices were incorporated into finished libraries, following manufacturer’s instructions. The final libraries were quantified by Qubit system (Thermo) and calibrated with an in-house control sequencing library. The size profiles of the pre-amplified cDNA and sequencing libraries were examined by Agilent Bioanalyzer 2100 using a High Sensitivity DNA chip (Agilent). Two indexed libraries were equimolarly pooled and sequenced on Illumina NOVAseq 6000 platform using the v2 Kit (Illumina, San Diego, CA) with a customized paired-end, dual indexing format according to the recommendation by 10x Genomics. Using proper cluster density, a coverage of around 250 M reads per sample (2000–5000 cells) were obtained corresponding to at least 50,000 reads/cell.

Twenty one biological samples (day 50; 4 WBS, 4 CTL, 4 7Dup) and (day 100; 3 WBS, 3 CTL, 3 7Dup) were examined by single-cell analysis with a coverage around 250 M reads per sample (2000–5000 cells) and at least 50,000 reads/cell as target. A total of 97,794 cells were retrieved after quality check with a minimum of 700 genes per cell and Mitochondrial RNA more than 5% per cell, to avoid low quality cells. Libraries from single-cell sequencing were aligned relying on the CellRanger v2.1 pipeline and using hg38 as reference.

Before downstream analyses, data deriving from the 21 samples was integrated by Conos (Barkas et al., 2019); the resulting clusters, common among all samples, were considered as shared populations and used as such for the scGen variational autoencoder algorithm (Lotfollahi et al., 2019). On the integrated dataset, UMAP dimensionality reduction as implemented in Scanpy (Wolf et al., 2018) was applied. Clusters were identified by applying the Leiden algorithm, a community detection algorithm that has been optimized to identify communities that are more coherent with the biological phenotype and more reliably identify cell populations. The resolution parameter value was optimized by surveying the stability of the resulting clusters. This resulted in the identification of final clusters.

Cluster annotation in cell populations was obtained by a combination of the following approaches: (I) scanpy’s rank_genes_groups to identify the most characterizing genes per clusters; (II) SCINA semi-supervised annotations algorithm (Zhang et al., 2019), using as source labels coming from three publicly-available datasets (López-Tobón et al., 2019; Nowakowski et al., 2017; Pollen et al., 2019), (III) Overlap of cluster marker genes (log2FC > 0.5, qval <0.05 compared to all other clusters) with cell type marker genes identified in a published single cell dataset (Nowakowski et al., 2017). Significance of overlap between marker sets was determined using Fisher’s exact test; (IV) projection of a single cell fetal cortex data in the UMAP by ingest algorithm. The obtained information was finally manually curated using a self-hosted cellxgene (chanzuckerberg/cellxgene: An interactive explorer for single-cell transcriptomics data (github.com), cellxgeneVIP (Li et al., 2020) and cellxgeneGateway (https://github.com/Novartis/cellxgene-gateway). Dotplots showing a panel of cell population specific genes across clusters was produced using the Scanpy DotPlot function.

Pseudotime analysis was performed on clusters identified by the Leiden algorithm with a resolution of 9. The resulting clusters were stratified by conditions/stage and then aggregated by their median to generate a new structure composed of ‘supercells’. Supercells with low contribution of reads for any genotype were discarded. The Palantir algorithm (Setty et al., 2019) was applied on control supercells (CTL) and then propagated.

For the neuron only cluster we applied the CellRank algorithm (Lange et al., 2022) using as starting point the lower pseudotime calculated in the whole population. We have recalculated the distance neighbor, the dimensionality reduction and al the subsequent calculation only on the neuronal cells, calculated the transition matrix for every genotype separated considering pseudotime (Setty et al., 2019), RNA velocity (La Manno et al., 2018) and similarity. We identify the terminal state using the CFLARE (Clustering and Filtering Left and Right Eigenvectors) to identify the eigenvalue and eigenvector pairs that are more probable and infer the terminal states. For The comparison with sh datasets and DUP we downscale the number of cells for DUP to be comparable in numbers, process not needed for the other conditions given the already comparable numbers.

For the comparisons of interest, the distribution of cells across clusters was analyzed by differential abundance analysis, performed following the procedure described in (http://bioconductor.org/books/release/OSCA/): edgeR was employed to test for differential abundance across genotypes A threshold of conventional p Value <0.05 was imposed to select significant modulations.

For Panel 3F we used Milo (Dann et al., 2022) and plot the differences between genotypes in the mature neuron clusters with p Value <0.05.

### Overlap with fetal marker genes

To refine the annotation of the sub-regions relative to the neural maturity clusters, we performed an overlap of the marker genes with the ones coming from a fetal dataset (Bhaduri et al., 2020), choosing GW22 sample. Fetal dataset was downloaded from UCSC cell data browser and the signatures were derived both from the reanalyzed data and the article.

The top-50 markers were then retrieved for each fetal cluster, as defined by the original annotation (Bhaduri et al., 2020); those not having top-50 significant marker genes (KS_PFC; KS_Outlier) or not of interest for the comparison with brain organoid were not considered in the overlap. Similarly, the top-50 marker genes were retrieved for the sub-regions of interest in the brain organoid neural maturity cluster, considering only the more differentiated clusters, identified by the external position in the force directed graph and the Cellrank algorithm (Lange et al., 2022); sub-clusters 1 and 6, not having 50 significant marker genes, were excluded from the overlap. Overlap analysis was performed with the multintersect function of the overlapper R package (https://github.com/plger/overlapper/) comparing the organoid and fetal top-marker lists and considering as the universe the genes used in brain organoid single cell analysis. The results are reported by a dot-plot[18] showing the number of overlapping genes; the dot is shown for overlaps having a PValue < 0.01; dot size shows the PValue and dot color the enrichment.

### Animals

All animal experiments were done in accordance with the Italian Laws (D.L.vo 116/92 and following additions), which enforces EU 86/609 Directive (Council Directive 86/609/EEC of 24 November 1986 and were approved by institutional ethics committee (OPBA committee) and the Italian Ministry of Health (Authorization 1073/16-PR).

Mice with hemizygous deletion (Gtf2i^+/-^) or duplication (Gtf2i^+/Dup^) of Gtf2i were generated as previously described (Mervis et al., 2012) and were maintained in an outbred CD-1 background. Mice were housed in groups of 5 max. in a normal 12 hr light:12 hr dark cycle and food and water were provided *ad libitum*.

### *In utero* electroporation

In utero electroporation was carried out as described (Saito, 2006). Timed pregnant mice were anesthetized using Avertin (1.25% solution, 0.02 ml/gr bodyweight) and the uterine horns exposed through a laparotomy. One microliter of endo-free purified (2 ug/ul) pCAG-EGFP plasmid (Addgene, cat# 89684) mixed with 0.05% Fast Green (Sigma, cat #F7252) in phosphate-buffered saline (PBS) was injected through the uterine wall into the lateral ventricle of the embryos. The DNA-injected embryos were held through the uterus, parallel to the embryonic anteroposterior axis. Then, five 40-V pulses of 50-ms duration at 1-s intervals were delivered across the embryonic head using 3-mm-diameter electrodes (Harvard apparatus, cat# 45-0487) connected to a square wave electroporator (Harvard apparatus, cat# EC1 45-0052). The uterine horns were placed back and the laparotomy was closed with sutures. The embryonic development could continue until E17.5 or postnatal (P) day 7.5. Following the surgery, Carprofen (0.5 mg/ml, 5 mg/kg bodyweight) was administered subcutaneously for the next two post-operative days.

### LSD1 inhibitor administration

The LSD1 inhibitor (*N*-[4-[(1*S*,2*R*)-2-aminocyclopropyl]phenyl]-4-(4-methylpiperazin-1-yl)benzamide ;dihydrochloride) was manufactured in the house by the drug discovery unit and was dissolved in 60% 5% glucose and 40% polyethylene glycol (PEG). Each mouse received 4 administrations of 10mg/kg LSD1 inhibitor or vehicle 2 times/week via oral gavage or 1 administration of LSD1 inhibitor or vehicle only for the bulk transcriptomic profiling experiment. Three hours following the last administration the mouse performed the behavioral test.

### Bulk transcriptomic analysis of mouse cortex

RNAseq FASTQ data were quantified at the gene level using Salmon (version 1.3.0). Mouse Genecode M25 was used as reference for quantification and annotation. 5 biological replicates were analyzed for each of the considered biological conditions (LSD1-inhibitor and vehicle, somatosensory and pre-frontal cortex, acute and chronic treatment). One replicate for the pre-frontal cortex chronic LSD1-inhibitor treatment was excluded from downstream analysis because identified as outlier in from quality control evaluations (sample-to-sample correlation). A total of 39 samples were included in differential expression analysis.

Protein-coding non-predicted genes with expression levels higher than 2 cpm in at least 4 samples were tested for differential expression by edgeR. Differential expression analysis comparing treated versus untreated mice was implemented separately for acute and chronic treatment, specifying the brain area as co-variable in the statistical model. Significantly modulated genes were selected setting a false discovery rate (FDR) lower than 10%. Analyses were performed in R, version 4.0.3.

### Behavioral experiments

Three-chamber sociability test was done as previously described (Moy et al., 2004). Mice 2-4 months of age were acclimatized to the experimental room for at least 1hr before the test. The test apparatus consisted of a 3-compartment box of transparent PVC with each compartment 20×40×22(h)cm (Ugo Basile). The test mouse habituated to the apparatus for 5 min. In the social preference session, the mouse could spend time either in the compartment with an empty metal grid enclosure (object) or in the compartment with a conspecific for 10 min. In the subsequent social novelty session, the mouse could spend time either with the familiar mouse from the previous session or with a novel mouse for 10 min. The apparatus was cleaned thoroughly with 70% ethanol and the compartments were alternated between mice. The data were acquired and analyzed using Any-maze behavioral tracking software (Stoelting). In the “washout” experiments, treated mice were left undisturbed in their home cages for 2 weeks and subsequently performed again the behavioral test as described above.

### Collection of biological materials

Blood samples were collected via the tail before drug administration and at the end of the behavioral experiments. In the “washout” experiments, an additional blood sample was collected at the end of the 4^th^ administration.

Following the behavioral experiments, the mice were sacrificed by cervical dislocation, the brain was rapidly removed and hippocampus, prefrontal and somatosensory cortices were dissected and snap-frozen in dry ice.

### Immunofluorescence

#### Cortical organoids

Cortical organoids were harvested at the indicated timepoint, fixed overnight at 4 °C in paraformaldehyde 4% PBS solution (SantaCruz). After rinsing with PBS, organoids were embedded in 2% low melting agarose dissolved in PBS; upon agarose solidification, blocks were put in 70% ethanol and kept at 4 °C before paraffin embedding, sectioning, and routine hematoxylin/eosin staining. Deparaffinization and rehydration were achieved by consecutive passages of 5 minutes each in the following solutions: 2 x histolemon (Carlo Erba), 100% ethanol, 95% ethanol, 80% ethanol and water. Sections were then incubated for 45 min at 95 °C in 10 mM Sodium citrate (Normapur)/ 0,05% Tween 20 (Sigma) buffer for simultaneous antigen retrieval and permeabilization; then left to cool for at least 2 hours at RT. To immunolabel the markers of interest, a blocking solution made of 5% donkey serum (ImmunoResearch) in PBS was applied for 30 minutes to the slides, while primary antibodies diluted in blocking solution were subsequently added, performing overnight incubation at 4 °C. Secondary antibodies and DAPI were diluted in PBS and applied to the sections for 1 hour and 5 minutes respectively. After each incubation step, 3 x 5 minutes of washing steps with PBS buffer were performed. After a final rinse in deionized water, slides were air-dried and mounted using Mowiol mounting medium.

Image analysis was done in ImageJ and the quantification was done by counting the number of positive cells relative to total DAPI positive nuclei.

#### Animals

At the corresponding developmental stages, the whole head (E17.5) or extracted brains (P7.5) were collected, immersed in 4% paraformaldehyde/PBS at 4°C for 4 h to overnight for fixation. Brains were washed twice with PBS, embedded into 3% low melting point agarose and then sectioned into 50-60 um using a vibratome (Leica, cat# VT1000S), mounted for direct visualization of EGFP or processed for immunofluorescence as it follows. Brain sections were washed briefly in PBS and blocked in PBS+ (PBS, 5% normal donkey serum, 0.25 % Triton-X100) for 2 h at room temperature. Sections were incubated with primary antibodies diluted in PBS+ overnight at 4°C. Following, three 5 min washes in PBS, secondary antibody diluted in PBS+ was added for 1h at room temperature (1:500). Sections were then washed 3 x 5 min in PBS and incubated for 5 min in DAPI (Invitrogen, cat# D1306) 1:1000 in PBS. After another 5 min wash in PBS, sections were mounted onto glass slides, coverslipped, and imaged using a confocal microscope (Leica SP8).

For image analysis and quantification of proliferative and neuronal markers we used custom made scripts in ImageJ. The choice of quantification and normalization method was based on manual inspection of image quality and/or marker expression. Specifically, for Phh3, Ki-67 and Bcl11b we quantified the total area of positive neurons and normalized it against the area covered by DAPI positive neurons. On the other hand, for Tbr2 and Cux1 we quantified the average intensity of the positive cells and normalized it against the average intensity of the DAPI positive neurons.

### Analysis of neuronal morphology

Morphology of entire neurons was visualized by the expression of pCAG-EGFP plasmid. Low cell-density dorsal telencephalic areas were used to determine morphological characteristics. Neurons transfected with plasmid DNA were selected by GFP expression and images were acquired at 60x magnification using a Leica SP8 confocal microscope. GFP-positive cells were traced with the NeuronJ plugin in ImageJ (FIJI package) for analysis of neuronal dendrite morphology following the developers’ instructions (Meijering et al., 2004). After the neurites were traced, they were labeled as primary (emanating directly from the soma), secondary (branching from a primary) or tertiary (branching from a secondary) and a text file was generated containing measurements of the lengths of all the neurites. The Snapshot tool in NeuronJ was used to save the tracings as an image file and these images were analyzed with the Sholl analysis plugin in ImageJ. The range of measurement was set using the straight-line tool traced from the center of the soma to the outermost neurite. Dendrite intersections were analyzed from a starting radius of 10 μm (to exclude the soma from analysis) with 5-μm steps to the outer radius. The resulting numbers of intersections per cell were used to calculate the mean and SEM for each radius interval.

## Author Contributions

A.L-T, R.S, G.T. Conceptualization, designed the experiments and wrote the manuscript. A.L-T, S.T, N.C. Developed organoids differentiation protocol. A.L-T, S.T, N.C, F.T, A.S, R.S. Organoids generation and maintenance. C.E.V and C.C. Developed data analysis pipeline, analyzed respectively single cell and bulk transcriptomic data and wrote the manuscript. R.S. Behavioral experiments and data analysis. P.F. Performed *in utero* electroporation and immunostainings in mice. E.T. Cell culture and libraries preparation. M.M. Sample preparation for proteomics. W.T.B. Provided fibroblasts. A.C, T.B. Proteomic analysis of organoids. C.M. and M.V. Designed and manufactured LSD1 inhibitor. L.O. Provided Gtf2i mice. G.T. Supervised the work.

## Acknowledgements

EPIGEN Flagship Project of the Italian National Research Council (CNR) (to G.T); the European Research Council (ERC DISEASEAVATARS no.616441 to G.T.; ERC PoC 713652–LSDiASD to G.T.); EDC-MixRisk, European Union’s Horizon 2020 research and innovation programme (Grant No 634880. to G.T.); ENDpoiNTs, European Union’s Horizon 2020 research and innovation programme (Grant No 825759 to G.T.); Telethon (GGP14265 and GGP19226 to G.T. and GGP19295 to A.L.-T.); Fondazione Cariplo (2017-0886 to A.L.-T.); Fondazione Umberto Veronesi (to R.S.); Ministero della Salute, RC 2019 – ERANET NEURON RRC-2019-2366750 -ALTRUISM (to M.M.); ERA-NET NEURON (ADNPinMED to P.F-B.); AIRC Grant No. IG-2018-21834 and EPIC-XS (project No. 823839, funded by the Horizon 2020 programme of the European Union to T. B.); The Leverhulme Trust and The Horizon 2020 Innovative Training Network EpiSyStem (Marie Skłodowska-Curie Actions; to A.S.). We are grateful to Federical Pisati from IFOM tissue processing facility, to Elaine Tam (University of Toronto) for assistance with mouse genotyping, to IEO Genomic Unit team, to IEO Imaging Facility team and Cogentech’s Animal Facility.

---

**Figure S1.**
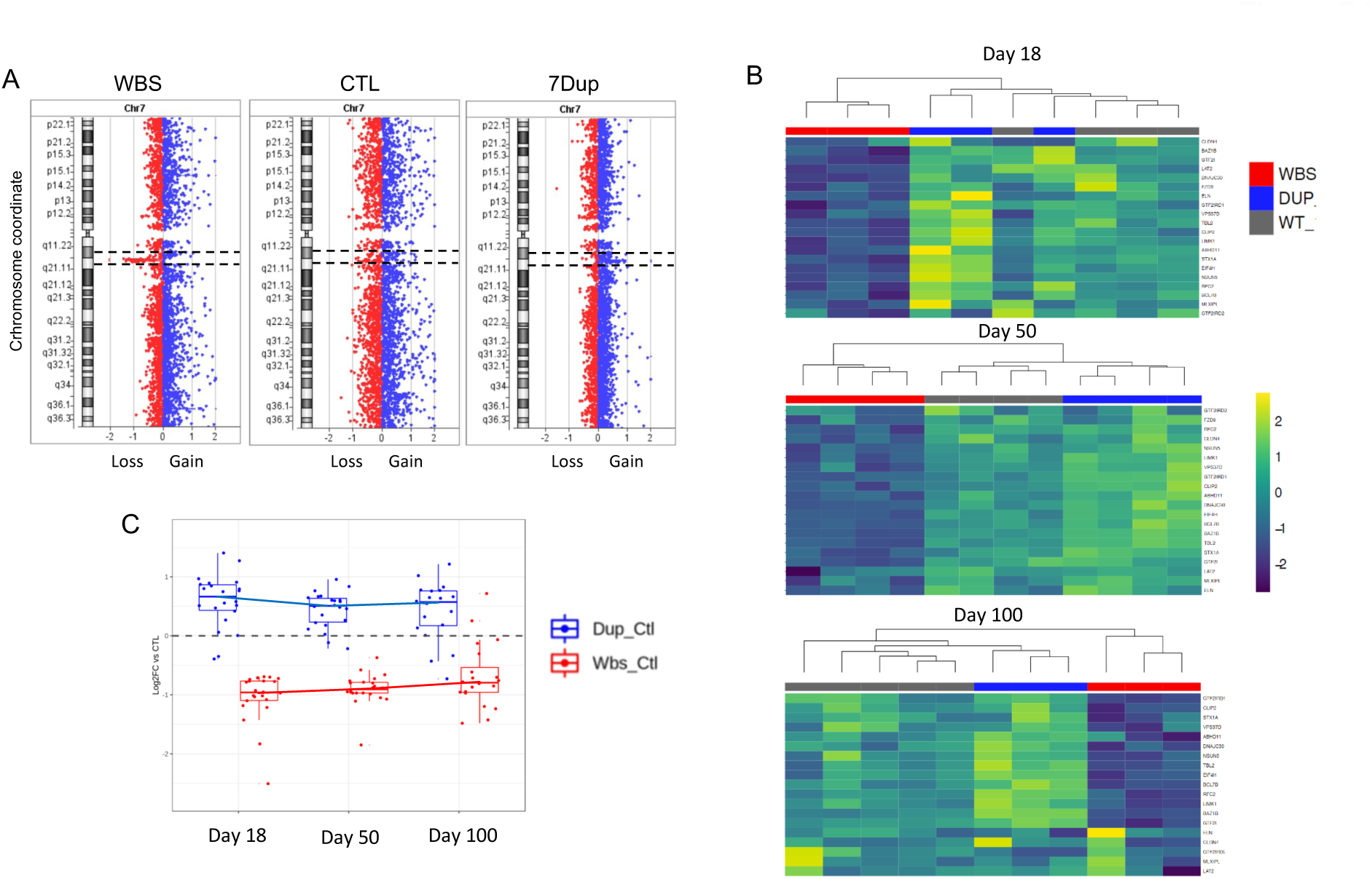
Expression levels of 7q11.23 genes in cortical brain organoids. **(A)** Array-CGH analysis for chromosome 7, representative of 1 iPSC line per genotype (WBS / CTL / 7DUP) dotted line highlights 7q11.23 locus. For details of the lines used in this study, refer to supplementary table 2. **(B)** Heatmaps reporting the expression levels of the genes located in the 7q11.23 CNV (z-score calculated on the Log2Cpm) from RNASeq transcriptome profiling in WBS, CTL and 7Dup cortical organoids at Day 18, Day 50 and Day 100 of differentiation. **(C)** Log2 fold-change of the genes located in the 7q11.23 CNV from stage-wise RNASeq differential expression analysis, comparing either WBS (in red) or 7Dup (in blue) to CTL. For each stage, the boxplot summarizes the values of each gene, visualized as a dot.

**Figure S2.**
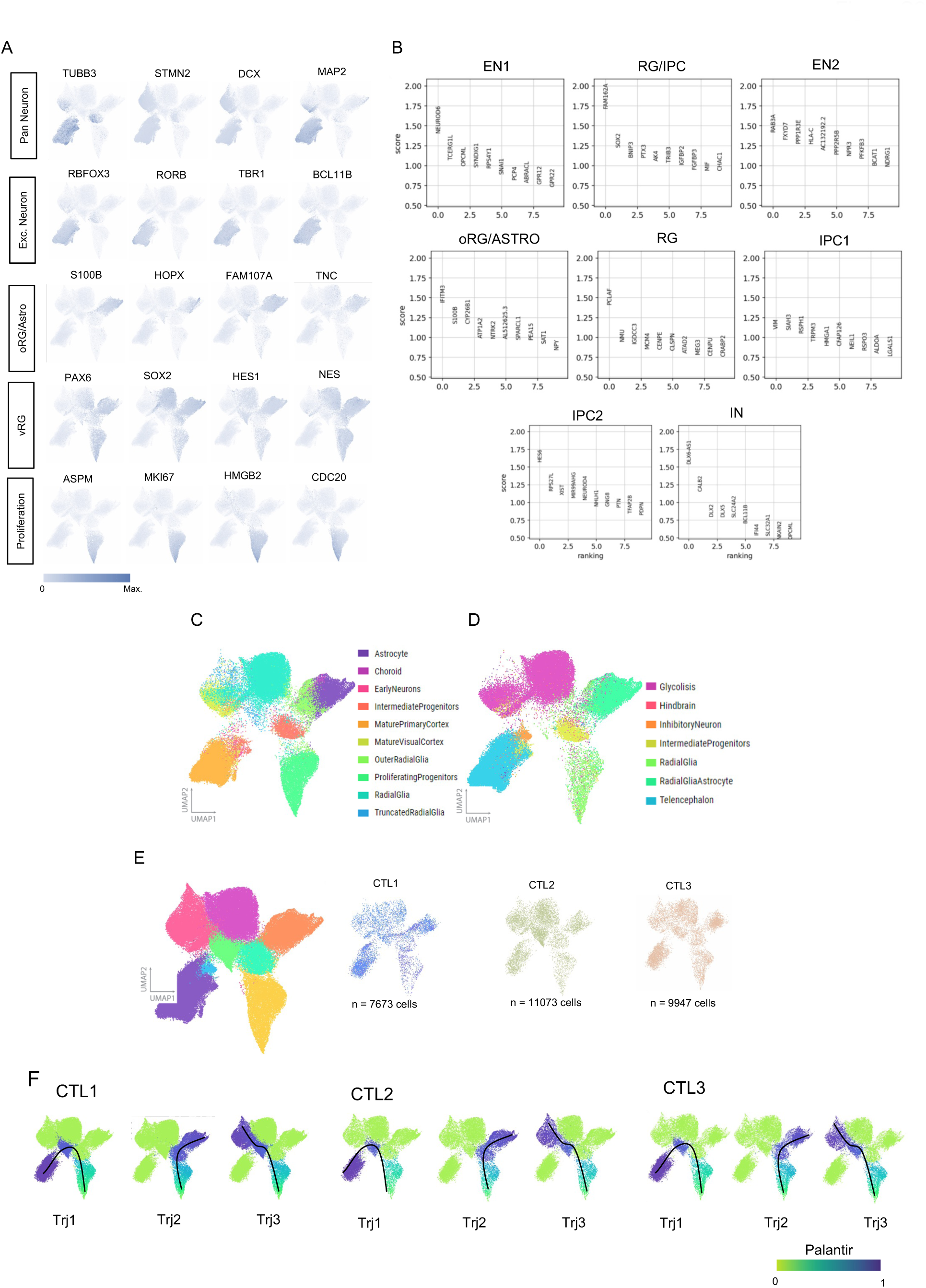
Cortical brain organoids show prototypical cell populations of cortex development. **(A)** Uniform manifold approximation and projection (UMAP) with color coded the most representative marker per population. **(B)** Top 10 marker genes for each identified cluster used for the manual annotations. **(C-D)** UMAP with color code representing automatically annotated cells using gene signature extracted from (Nowakowski et al., 2017; Pollen et al., 2019) using SCINA (Zhang et al., 2019). **(E)** UMAP with color code representing cell distribution in the UMAP among the 3 CTL lines. **(F)** UMAP with color code representing the 3 main differentiation trajectories, in all control genotypes are the same.

**Figure S3.**
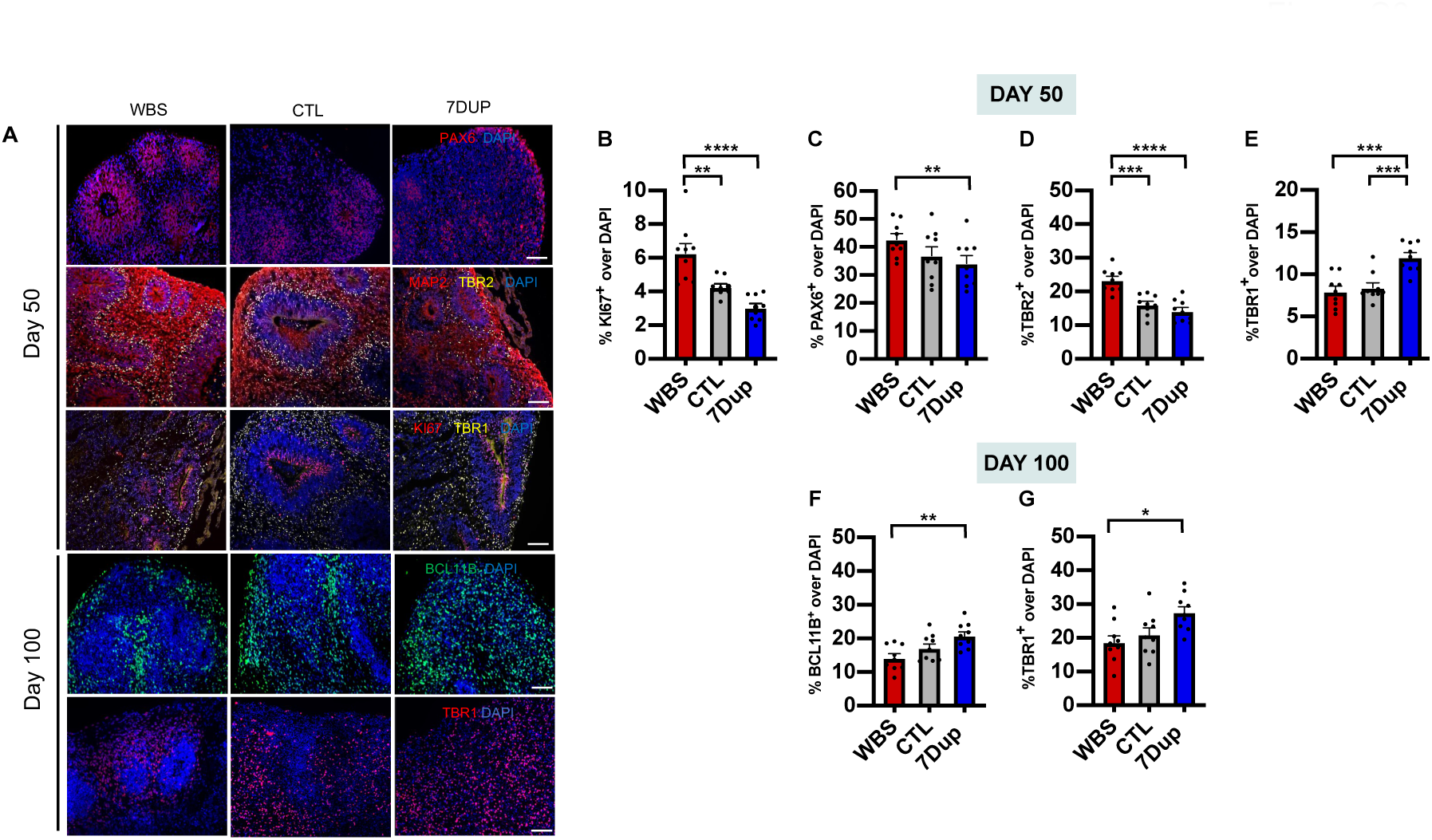
Accelerated neuronal differentiation dynamics in 7Dup COs. **(A)** Representative widefield fluorescence images from day 50 and day 100 cortical organoids across genotypes (WBS / CTL / 7Dup), immunostained with markers for different cortical populations: PAX6 (apical radial glia), MAP2 (Neurons) TBR2 (intermediate progenitors) TBR1 (Layer VI neurons) Ki67 (cycling cells) BCL11B (Layer V-V neurons). Scale bar, 50 um. **(B-E)** Quantification of the percentage cells positive for different population markers relative to total nuclei (DAPI) at day 50 of differentiation. **(F and G)** Quantification of the percentage Layer V and VI neuron markers relative to total nuclei (DAPI) at day 100 of differentiation. Each data point is an individual section from five organoids per line, three independent hPSC lines per genotype. All data are shown as mean ± SEM (27 sections from 3 COs/cell line, 3 cell lines/genotype), one-way ANOVA, post-hoc Tukey test; *p < 0.05, **p < 0.01, ***p < 0.001.

**Figure S4.**
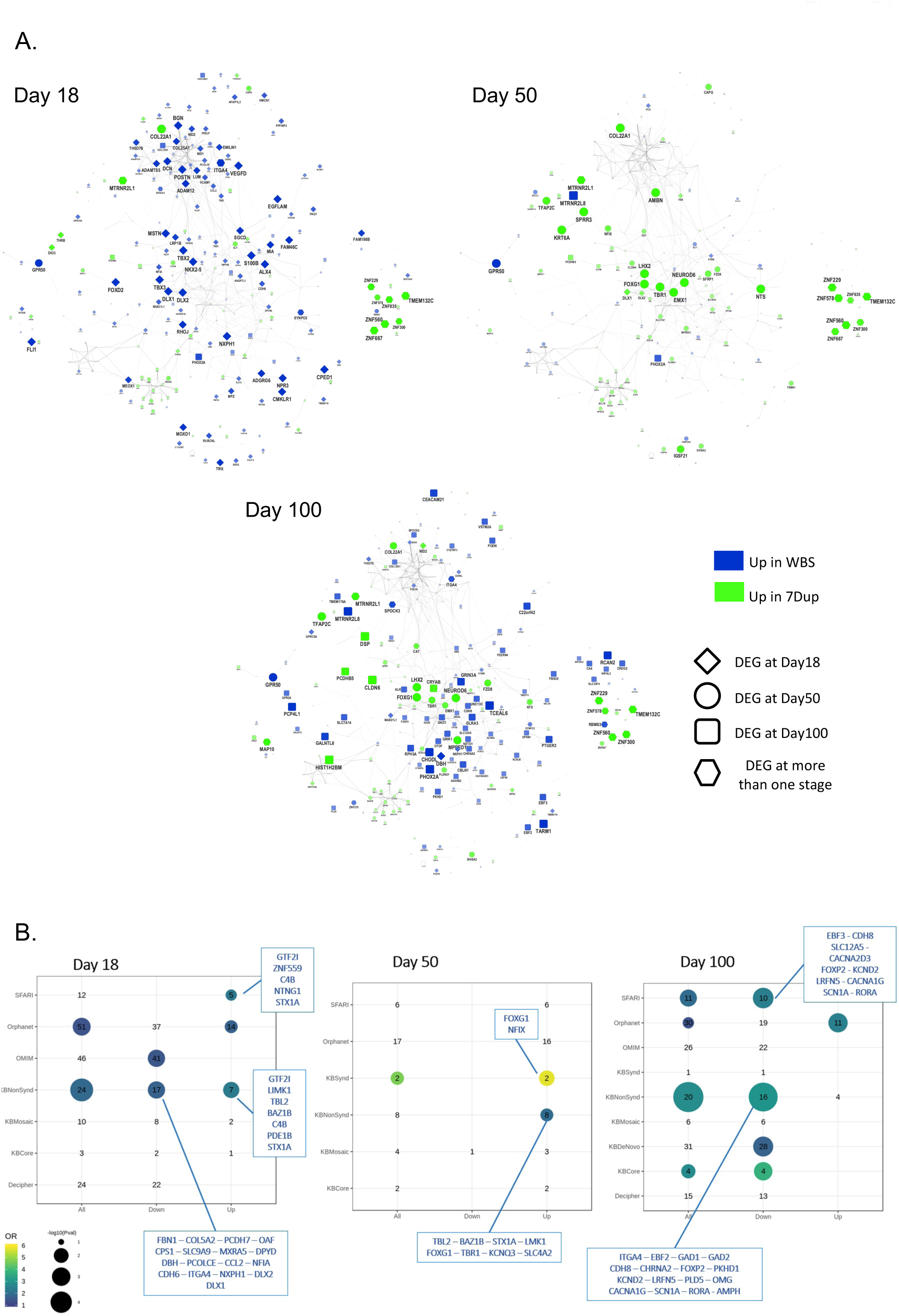
Genes modulated by 7q11.23 gene dosage imbalance significantly overlap with disease-related knowledge bases. **(A)** STRING-based (https://string-db.org/) network reconstructed for the genes found modulated (FDR < 0.1, log2FC > 1) in at least one time-point by stage-wise differential expression analysis. At each examined stage, node (gene) size and color represent respectively the magnitude and direction of Log2FC, with genes more expressed in 7Dup and WBS in green and blue respectively. Across the three visualizations, node shape highlights the stage in which the genes are significantly modulated: diamond for Day 18, circle for Day 50, square for Day100 and hexagon for genes significant in more than one stage. **(B)** Dotplot showing the results of the overlap analysis between genes differentially expressed in 7Dup compared to WBS and disease-related knowledge bases. DEGs (FDR < 0.1, log2FC > 1 as absolute value, stage-wise DEA in cortical organoids) are analyzed either grouped or split in up- and down- regulated. The plot reports the number of overlapping genes for Odds Ratio (OR) > 1 and the dot for p-values < 0.1. Dot size and color represent the p-value and the OR respectively.

**Figure S5.**
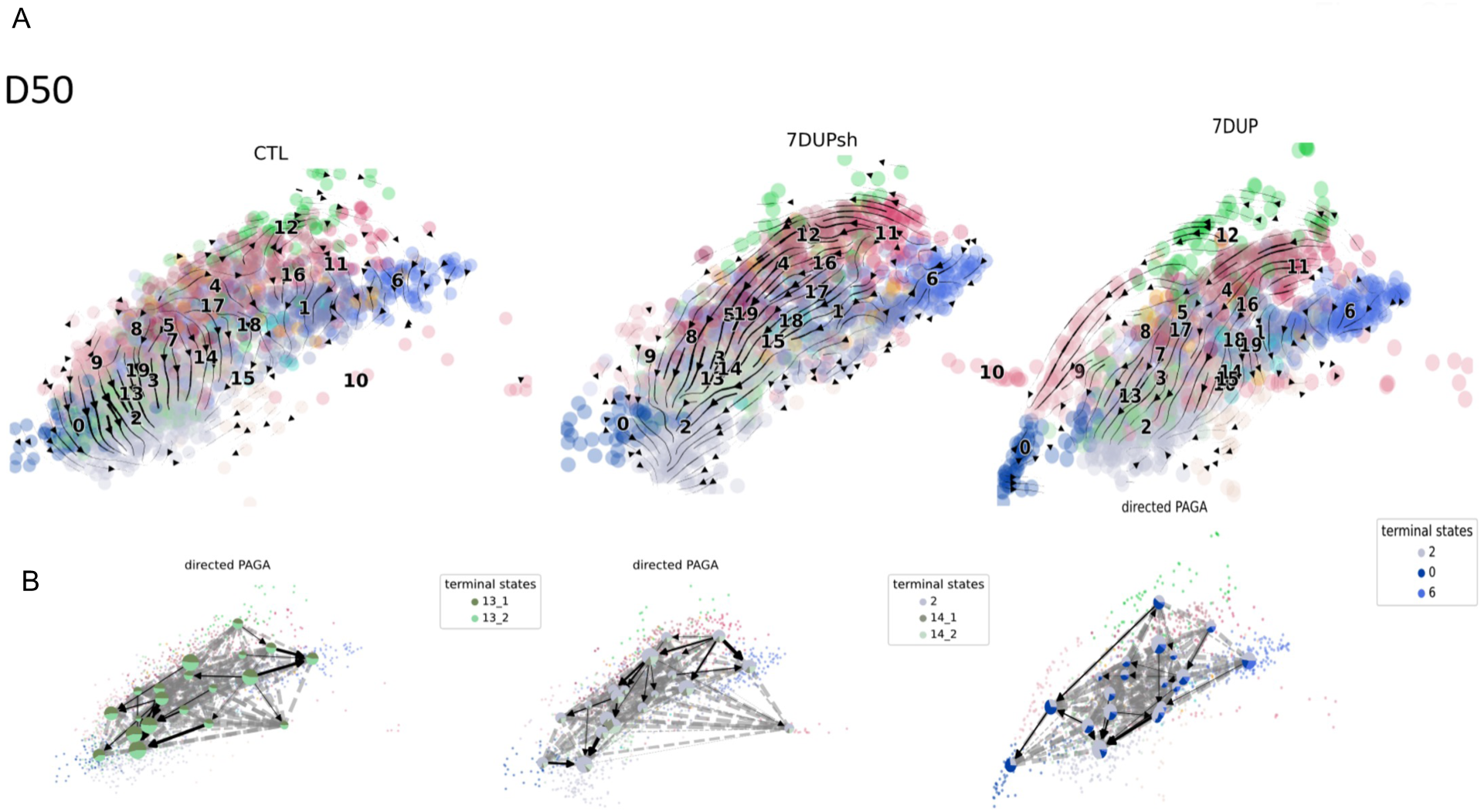
Gene dosage imbalances at 7q11.23 dysregulated normal differentiation trajectory at D50 and partial rescue by the knockdown of GTF2I. **(A)** Force-directed graph stratified for genotype (CTL, 7DupshGTF2I and 7Dup) from left to right at D50, with color coded the identified subpopulations. The arrow defines the inferred differentiation trajectory identified by Velocyto. **(B)** PAGA with position identified by force-directed graph subpopulations location. Color code identifies the number of cells per cluster that will be most probably going in the identified terminal state.

**Figure S6.**
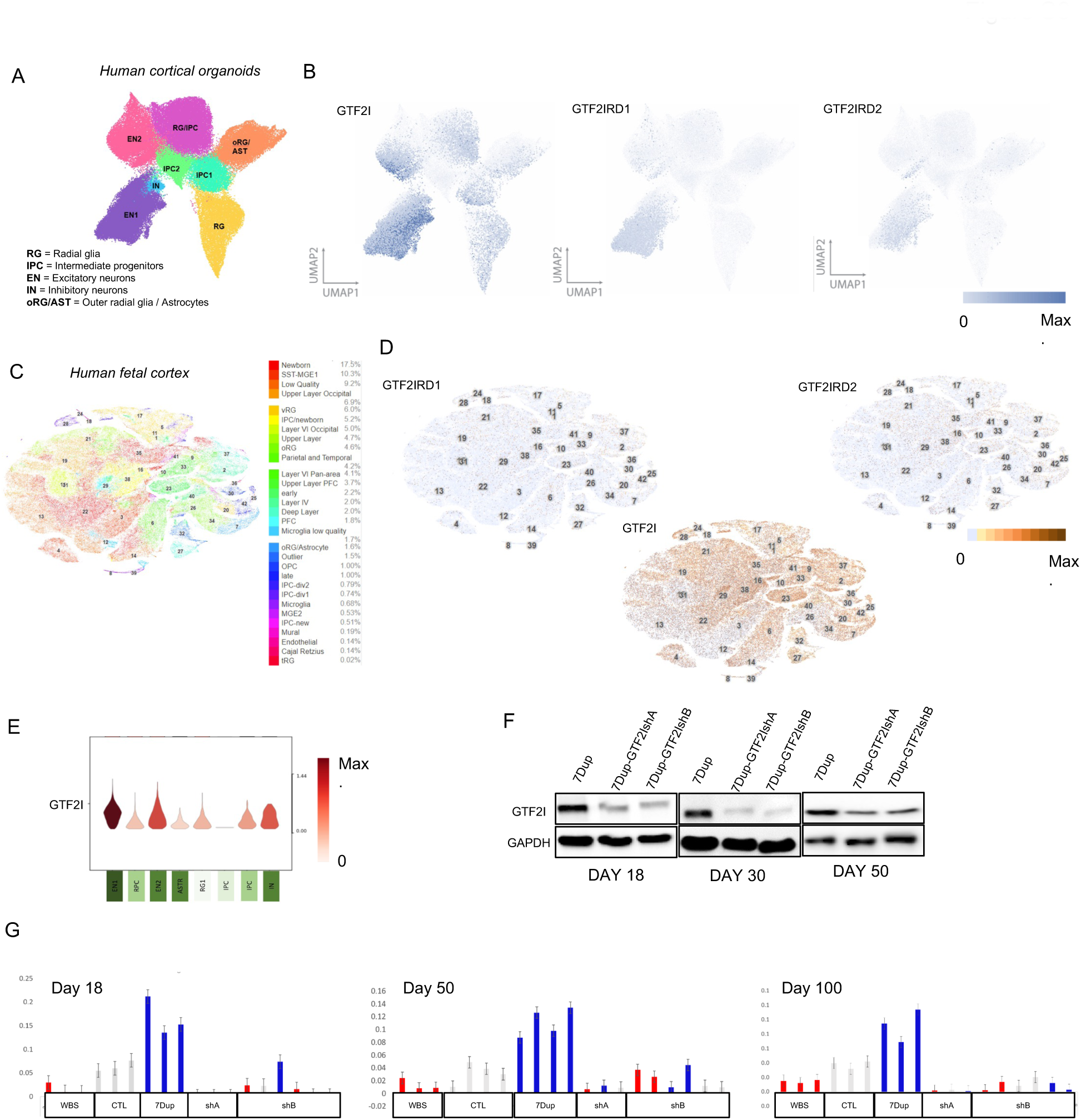
GTF2I presence in fetal brain and organoid depending of the cell populations. **(A)** UMAP with color coded the identified population. **(B)** UMAP showing expression levels in control COs for GTF*2I, GTF2IRD1* and *GTF2IRD2*. **(C)** Seurat tSNE showing the population annotation from nearly 200,000 primary cells sampled from GW6-22 from seven cortical regions, including PFC, V1, motor, somatosensory, temporal, parietal and hippocampus. Annotation taken from the UCSC cell data browser. Cell Browser dataset ID: organoidreportcard/primary10X. **(D)** Expression levels in human fetal primary brain cells for *GTF2I, GTF2IRD1* and *GTF2IRD2.* **(E)** Breakdown of expression levels in control in COs for GTF*2I* between different populations from a single cell dataset. **(F)** Western blot showing the downregulation of GTF2I using two different short hairpins respectively, at Day 18, 30 and 50, using GAPDH as loading control. Each lane is representative from a pool of n = 3 organoids/cell line, 2 cell lines/genotype. **(G)** Expression levels of GTF2I in COs at Days 18, 30 and 50 by quantitative RT-PCR. Values are expressed as normalized expression over the average for GAPDH levels. Each bar represents a pool of n = 3 organoids from a single organoid line.

**Figure S7.**
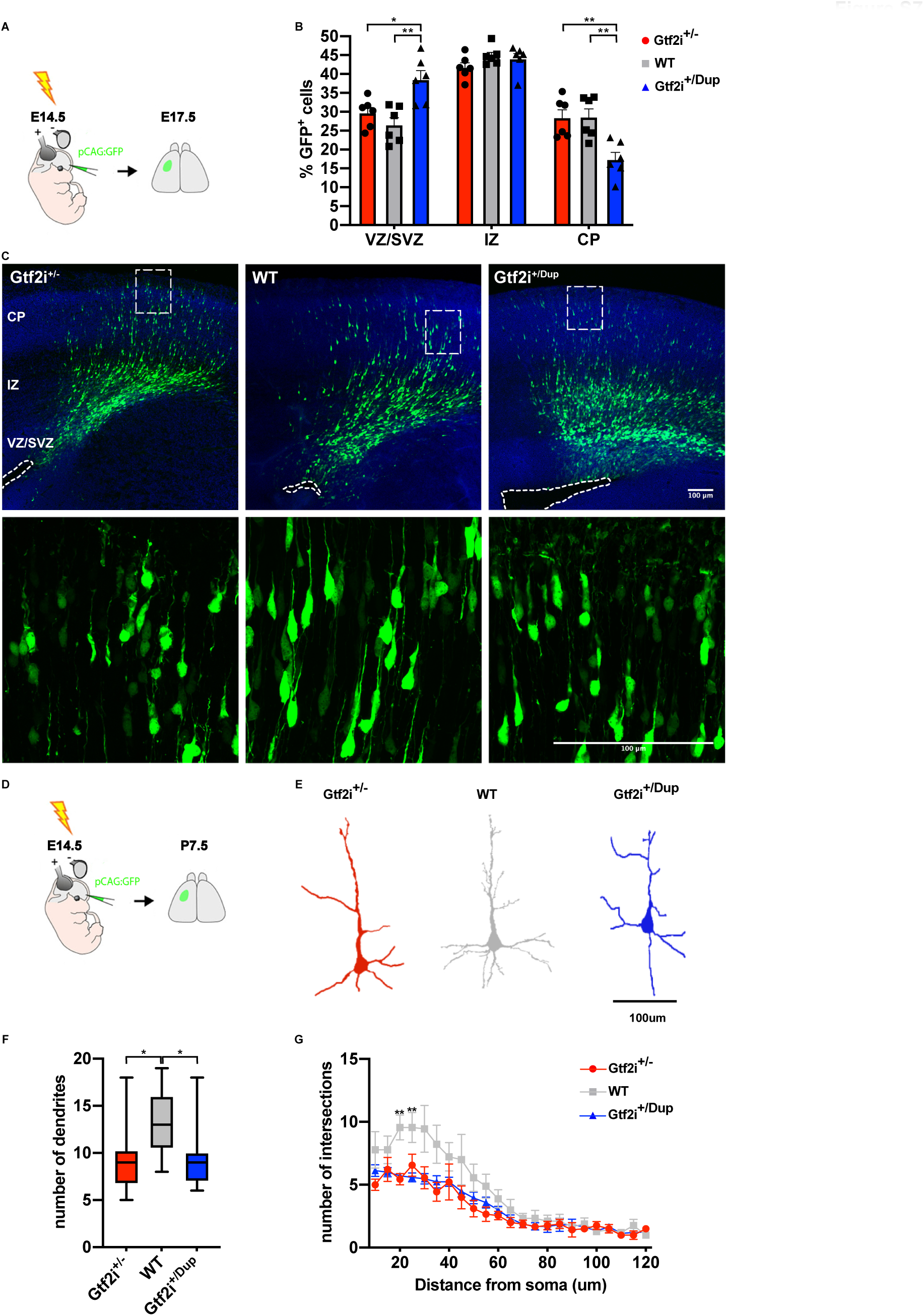
Gtf2i dosage affect neuronal migration and neuronal morphology in mouse cortex. **(A)** Schematic representation of *in utero* electroporation experiments. Mice were electroporated at E14.5 and analysed at E17.5. **(B)** Bar diagram depicting neuronal migration defects in Gtf2i^+/Dup^ mice. Note the difference between increased GFP^+^ neuronal progenitors in VZ/SVZ and reduced GFP^+^ neurons in CP. Gtf2i^+/-^, WT, Gtf2i^+/Dup^; n=6 sections from 3 mice/group. Data shown as mean±SEM. One-way ANOVA followed by Tukey’s multiple comparison test. Significance level was set to p<0.05. *p<0.05; **p<0.01. **(C)** (Upper panel) Representative images of electroporated brains from Gtf2i^+/-^, WT and Gtf2i^+/Dup^ mice at E17.5. (Lower panel) Insets of upper panel figures showing GFP^+^ neurons in CP. **(D)** Schematic representation of *in utero* electroporation experiments. Mice were electroporated at E14.5 and analyzed at P7.5. **(E)**. Representative morphology of cortical neurons from Gtf2i^+/-^, WT and Gtf2i^+/Dup^ mice at P7.5. **(F)** Box plot showing reduced number of dendrites in mutant mice compared to WT. Gtf2i^+/-^, n=10 neurons; WT, n=9; Gtf2i^+/Dup^, n=15. Kruskal-Wallis followed by Dunn’s multiple comparisons test. **(G)**. Plot showing number of intersections from Sholl analysis. Gtf2i^+/-^, n=19 neurons; WT, n=9; Gtf2i^+/Dup^, n=15. Data shown as mean±SEM. Kruskal-Wallis followed by Dunn’s multiple comparisons test. Significance level was set to p<0.05. *p<0.05; **p<0.01.

**Figure S8.**
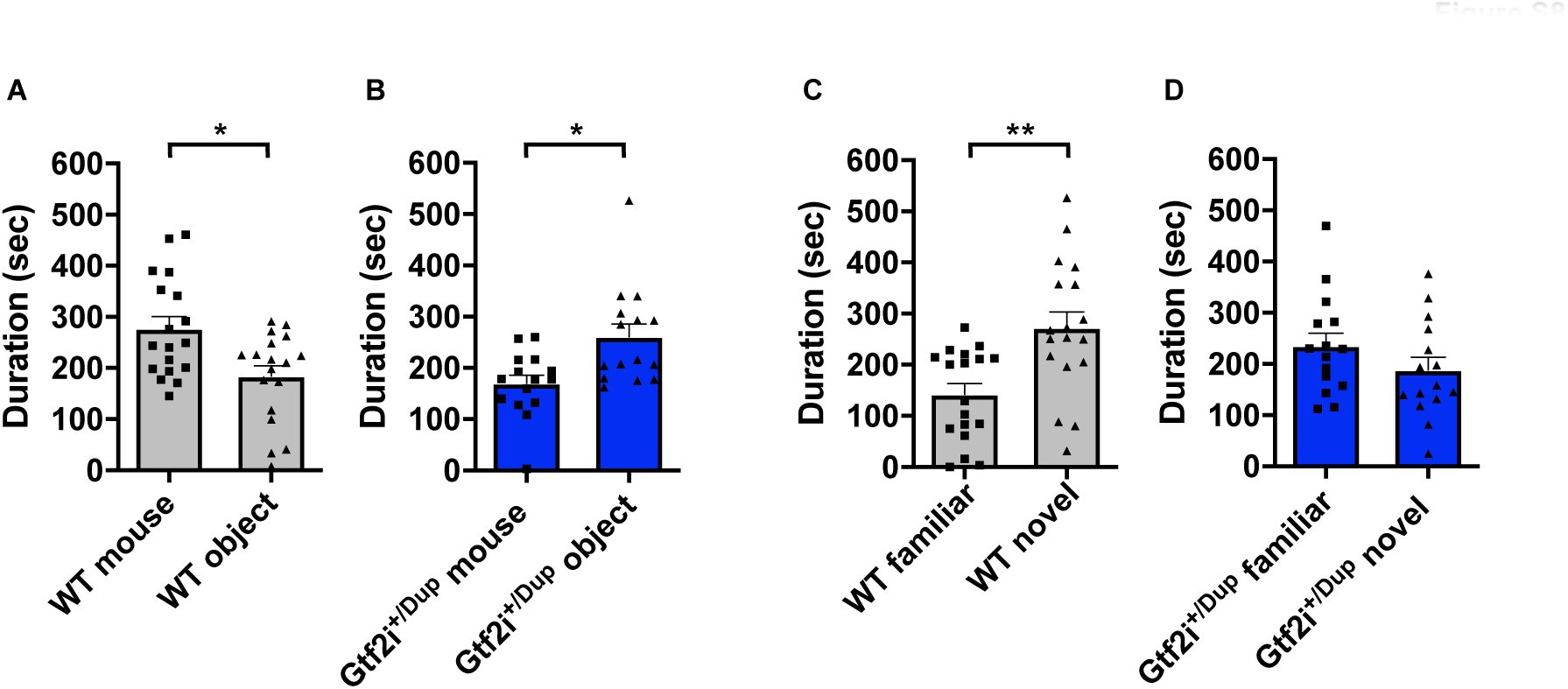
Gtf2i duplication causes social preference impairments in female mice. Bar plots with dots depicting the quantification of social preference **(A, B)** and social novelty **(C, D)** in female Wt (n=18) and Gtf2i^+/Dup^ mice (n=15). Statistical analyses were performed using paired Student’s t-test followed by Holm-Bonferroni correction for multiple testing. Significance level p<0.05. *p<0.05; **p<0.01.

**Figure S9.**
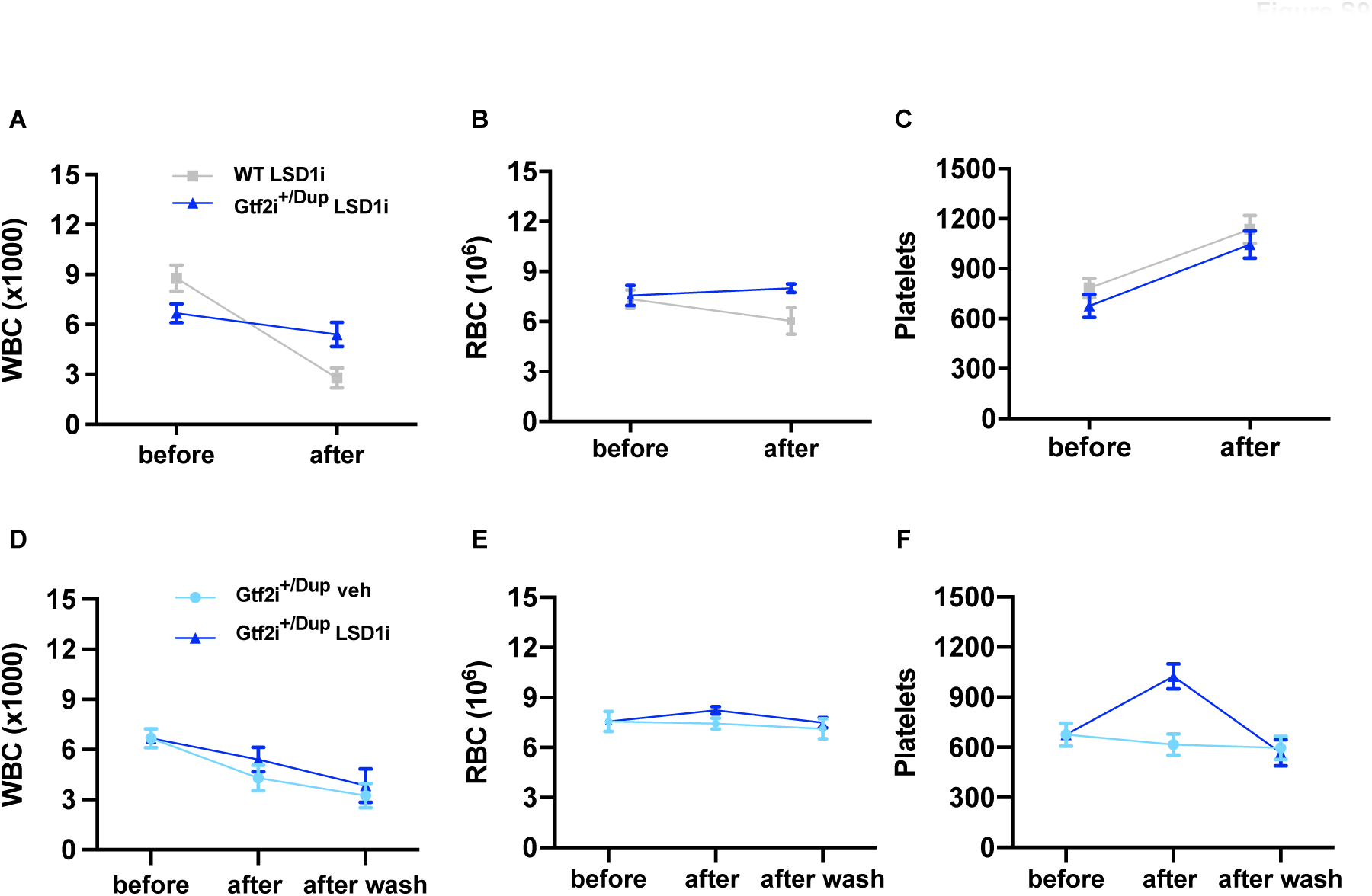
Blood factors remain within normal range following LSD1 inhibitor administration. **(A-C)** Line charts depicting values respectively, of white blood cell (WBC), red blood cells (RBC) and platelets before (Wt n=18; Gtf2i^+/Dup^ n=16), and after (Wt n=8; Gtf2i^+/Dup^ n=21) administration of LSD1 inhibitor. **(D-F)** Line charts depicting values respectively, of WBC, RBC and platelets in Gtf2i^+/Dup^ mice before (n=16), after administration of LSD1 inhibitor or vehicle (LSD1i n=21, vehicle; n=16), and after drug ‘washout’ (LSD1i; n=13; vehicle; n=11).

**Table S1.**
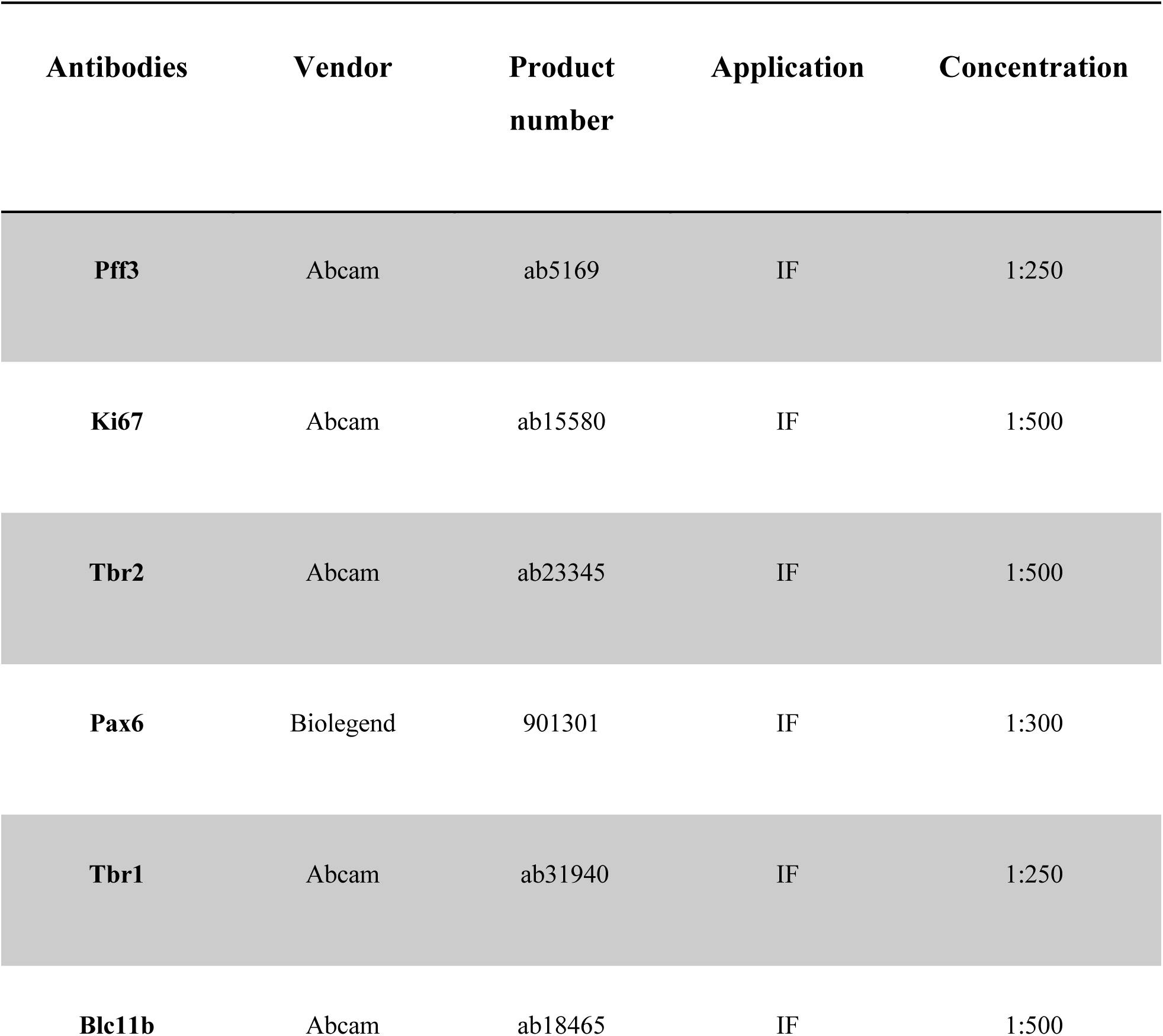
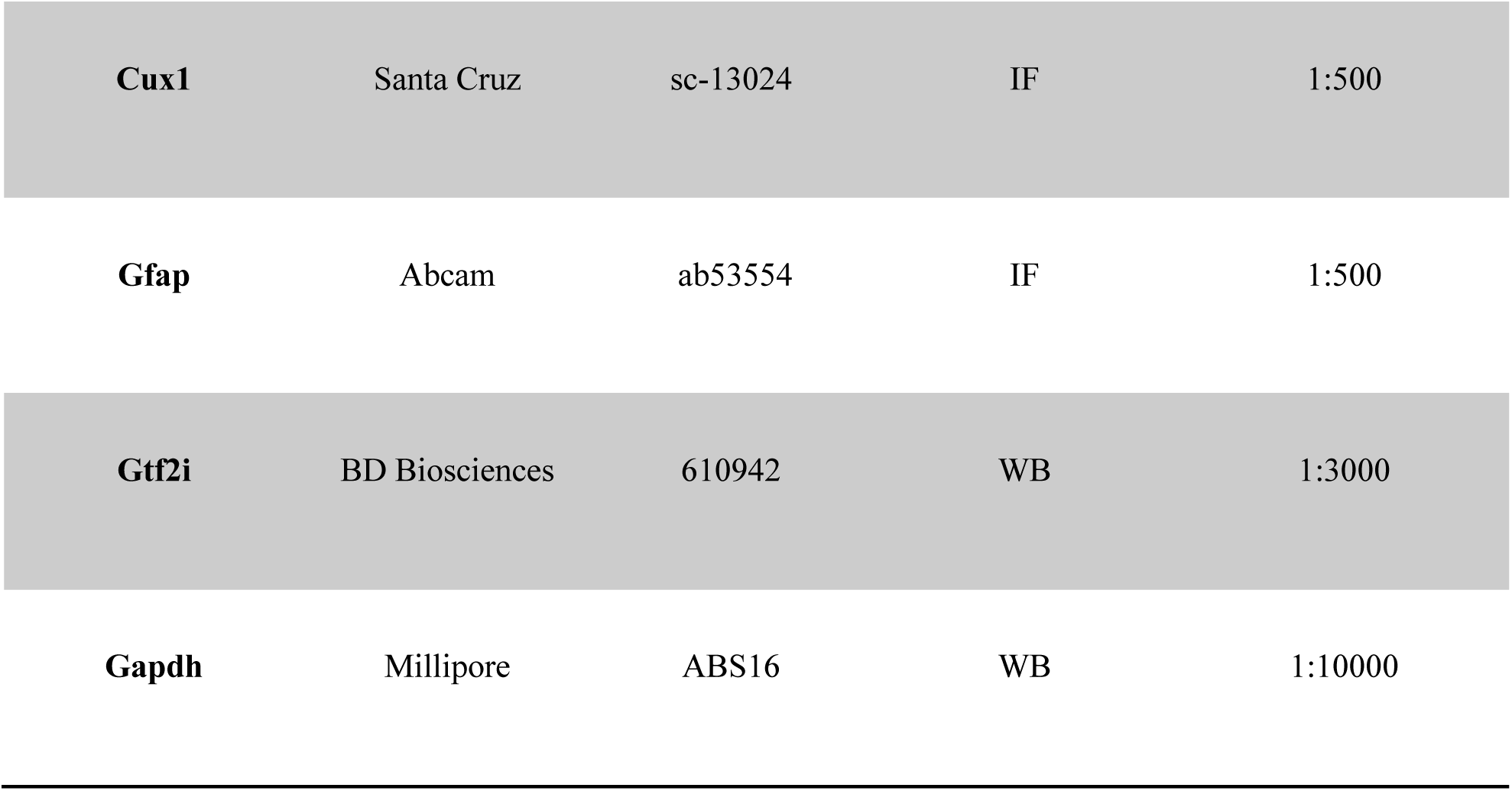
Antibodies dilutions

